# Humidity-driven shape morphing enhances fog harvesting in porous cactus spines

**DOI:** 10.1101/2025.10.07.676731

**Authors:** Jessica C. Huss, Finn Box, Qun Zhang, Martin A. Grömmer, Sebastian J. Antreich, Tofayel Ahammad Ovee, Jean-François Louf, Jürg Schönenberger, David G. Williams, Notburga Gierlinger, Mingchao Liu, Kevin R. Hultine

## Abstract

Cacti develop spines instead of conventional leaves, which often serve as mechanical defence against herbivores and protection from excess sunlight. However, some cactus species grow porous and flexible spines, suggesting fundamentally different functions. Here we demonstrate that the porous spines of *Turbinicarpus alonsoi*, a cactus native to central Mexico, function as hygro-morphing fog harvesters. We discovered that the spines are highly hygroscopic and straighten when exposed to fog, leading to increased fog water collection rates. Experiments and numerical simulations confirm that straightening is driven by swelling-induced pressure in the cell walls of the spine tissue. Swelling results from capillary imbibition of fog water and predominantly generates expansion in the transverse plane, which causes the pre-curved spines to straighten. Despite their porosity and hygroscopicity, the spines prevent direct absorption of fog water into the living cortex due to the presence of a suberin-rich tissue layer at the spine base that instead promotes surface runoff towards the roots. Our work suggests that hygro-morphing emerges from distinct structural, biochemical and geometric adaptations of cactus spines, and enables a fine modulation of the flow dynamics on the surface of spines. The increasing plant water supply from fog by shape morphing provides a potential advantage for survival in a hot, semi-arid region with frequent fog formation.

**Significance statement:** In many arid regions of the world, fog is a critical source of fresh water. Plants display striking ways to harvest this fog water. For cacti with porous spines, it has long been assumed that they collect fog water directly by capillary imbibition. However, we demonstrate that, in *Turbinicarpus alonsoi*, water-impermeable tissue at the spine base prevents direct transport of water into the living cortex. Instead, a dual process increases fog water collection: the curved spines initially imbibe fog water, which causes them to swell and straighten, and a thin liquid film then forms on the spine and runs off along the plant surface down to the roots. Hygro-morphing spines therefore enhance the capacity of cacti to collect fog water.

## Introduction

Living plant tissues regulate their turgor pressure to induce organ movement and to modulate shape (*1*–*3*), while non-living (dead) plant tissues often generate movement from swelling and shrinkage of hygroscopic cell walls (*4*–*6*). Retaining dead but functional tissues is thus an effective strategy for plants to decouple movements from metabolic processes and subject them to environmental control, such as changes in relative humidity or rainfall. These humidity-driven shape changes are known as hygro-morphing (*7*) and typically occur in connection with seed release and dispersal, as shown in seed pods of orchid trees (*8*), ice plants (*9*) and pine trees (*10*), as well as in awns of filaree and wheat (*11-13*), and in the pappus of dandelions (*14*). In these examples, movements typically arise from differential expansion of adjacent tissue layers in response to hydration or dehydration (*8-12, 14*). This heterogeneous mechanical response to water can result in large deformations, including the bending, coiling and twisting that is prominent in slender plant structures (*8, 11-13*). Shape-morphing plant tissues typically display at least two distinct layers that either differ in their internal orientation of fibres and fibrils (*6-8, 13*), or in their capacity to absorb water due to biochemical differences (*14*). As a result, the layers expand differently (in different directions and/or relative amount), leading to mismatches in mechanical strains that induce internal stresses and cause the overall structure to deform.

Although hygro-morphing is widespread in nature, we are unaware of it having been reported directly in the context of fog harvesting – specifically, in systems where fog itself triggers the hygro-morphing, which in turn enhances fog collection rates. Fog events are generally characterized by high ambient humidity, i.e. a relative humidity of RH > 95%, and serve as a potential, and sometimes critical source of fresh water for plant survival in arid regions (*15-17*). Favourable conditions for fog harvesting exist in zones with advection and orographic fog (*18*). In recent years, several engineered devices have been developed to optimize fog collection in such environments, including conventional Raschel and stainless-steel meshes (*19*), mesh-like kirigami structures (*20,21*), wire-based harps (*22*), as well as bio-inspired systems, such as sequoia leaf-inspired harp meshes (*19*) and cactus-inspired spine arrays with tuned surfaces (*23-26*). A notable increase in the aerodynamic collection efficiencies of planar collectors, such as meshes, can be achieved by introducing a curvature, which reduces the flow resistance of the mesh and maintains higher downstream velocities (*18, 27*).

Characteristic features of plants in fog-rich environments are long, narrow leaves and a rosette growth form, which facilitate fog interception, droplet coalescence and gravitational drainage (*15, 28*). Ultimately, most plants absorb the collected fog water by their roots; except for certain species of *Tillandsia*, which lack a proper root system and possess specialized trichomes on the leaf surface for direct absorption of fog (or dew) droplets (*16*). Similarly, it has been suggested that some species of cacti also absorb fog droplets directly via their spines (*29-33*). However, this has not been investigated systematically so far, even though many cactus species grow in arid and semi-arid regions with frequent fog formation and produce porous spines, which could suggest the existence of such a mechanism.

The western parts of the Sierra Gorda mountain range in central Mexico are ideal to explore fog harvesting mechanisms in detail, because this region is characterized by nocturnal fog events and montane cloud belts (*28, 34*), as well as high species richness of cacti (*35*), especially in the genus *Turbinicarpus* (*36*), known for its porous spines (*31, 37*). Consequently, in this work, we studied the porous spines of *Turbinicarpus alonsoi* – a critically endangered cactus species (IUCN Red List, 2009) endemic to a small mountainous, fog-rich region with a hot, semi-arid climate – and demonstrate that the spines imbibe fog water but inhibit direct absorption into the living plant cortex. Instead, water is transported superficially towards the root system for absorption underground. Intriguingly, we also discovered that the highly hygroscopic spines straighten when exposed to a stream of fog, which increases fog water collection rates. The highly adapted spines offer unique insights into the functional diversification of cactus spines and may serve as botanical inspiration for engineered systems, particularly in the context of fog harvesting (*19, 22*), shape morphing and autonomous materials (*38-41*).

## Results

### Fog exposure leads to spine straightening

At maturity, the spines of *T. alonsoi* solely consist of dead tissue and are strongly curved, which can be achieved by differential growth (*42, 43*). When the dry and curved spines are exposed to a stream of fog at low velocity (avoiding any drag-induced reconfigurations), they absorb fog water droplets immediately and steadily straighten over the course of minutes (Fig. 1A; Movie S1). Each spine deforms differently; longer spines from the apical region of the plants predominantly show longitudinal straightening, leading to an increase in spine height and a more vertical orientation when exposed to fog (Fig. 1B-C). From 2D images, the deformation of individual spines can be quantified by measuring the height *H*_*t*_ of a spine of length *L* (in the dry state) as a function of time *t* (Fig. 1B). During constant frontal fog exposure, *H*_*t*_*/L* initially increases linearly before saturating around a constant value (Fig. 1C). To test how changes in spine inclination influence the amount of collected fog water, we varied the base angle *α* of individual spines (with similar lengths) to generate different initial spine heights and exposed them to a laminar stream of fog at constant intensity for *t*=1800 s repeatedly (Movie S2, Fig. 1D). We found that the amount of fog water collected by the spines is linearly proportional to the final spine height (*H*_*1800*_*/L*) in the tested range of 0.6 < *H*_*1800*_*/L* < 1.0. This demonstrates that hygro-morphing of spines increases fog-water collection rates. Bending across the thickness results in an increase in spine height and alignment with gravity (Fig. 1), which promotes gravitational drainage (*44*), see also Fig. S1A). Some spines were also observed to twist as they uncurled; however, twisting is a secondary effect – the shape of the cross-section renders spines less resistant to bending normal to their thinnest surface (dorsiflexion) and more resistant to torsion, i.e. twisting (*45, 46*). Although twisting can increase the distance between individual spines on the same areole (Fig. 1A), which may facilitate direct exposure to fog and thereby enable more efficient droplet collection from various directions (*41*), here we focus on bending since it is the dominant mechanism for shape-morphing that leads to uncurling and increases spine height.

**Fig. 1.**
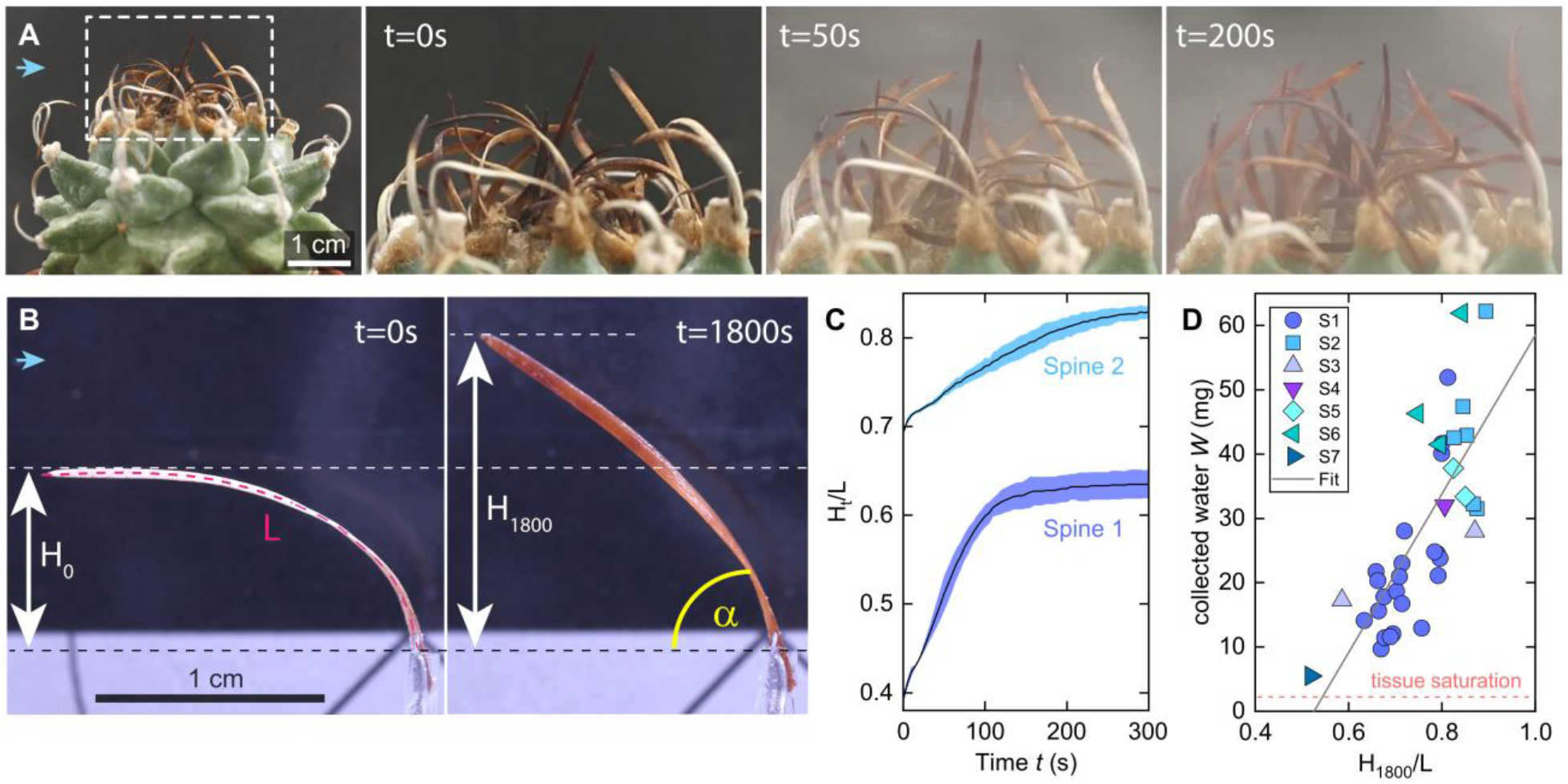
Spine straightening and water collection during fog exposure in *Turbinicarpus alonsoi*. **(A)** The spines are arranged in a rosette-like manner on the mature plant with longer spines growing in the apical (top) region. When exposed to fog (flow direction indicated by blue arrow), the initially dry and curved spines (*t=0 s*) steadily deform (*t=50 s*) and straighten up (*t=200 s*). **(B)**A single spine with length *L* shown in the dry state (*t=0 s*), with height *H*_*t*_ indicated before (*H*_*0*_) and after *t=1800 s* of fogging (*H*_*1800*_, in the wet state), with fog incoming from the left. The angle *α* at the spine base increases during straightening, along with spine height. (**C**) *H*_*t*_*/L* measured over time during the initial phase of fog exposure for spine 1 (shown in B) and spine 2 from the same plant (means ± SD with *n*=3 repetitions for each spine), showing different height gains during fogging. (**D**) Total amount of fog water (*W)* collected by the spine tissue of *n=7* different spines and the glass capillary, in which the base of the spine is inserted, during a 30-minute fogging period, shown as a function of *H*_*1800*_*/L*. Each data point (*n=37*) represents the total amount of fog water that a single spine collected during fogging; the initial spine height *H*_*0*_ was varied systematically by changing the angle α at the base of the spine (shown in panel B). The mean saturation limit of the tissue is indicated, as well as a linear fit (*W*=-65+124(*H*_*1800*_*/L*)) applied to all data points.

A small amount of water is required to initiate shape-morphing and is bound within the spine tissue. In our experiments, this was apparent from the delayed onset of water accumulation in the glass capillary attached to the base of the spine, which occurred only after the spine tissue had already absorbed an initial amount of fog water (Fig. S1A). Since the exact amount of water that the spines absorbed during fogging could not be measured during experiments, we immersed the tested spines in liquid water for *t*=30 min (1800 s) to determine the amount of water stored in saturated spines. Saturated spines absorbed *m*=0.74 ± 0.22mg (mean±SD of *n*=10) of water per milligram dry mass, which corresponds to a relatively small share of the total amount of fog water collected by the spines (mean tissue saturation *W*_*s*_=2.25 ± 0.22 mg in Fig. 1D; ±SD of n=10) and indicates that the bulk of the collected water drains off the spine surface.

### Dynamics of water imbibition and drainage

The interaction of the spine tissue with fog droplets is characterized by two consecutive, but temporally overlapping processes: (i) capillary imbibition occurs via surface cracks and is promoted by the exposed hydrophilic internal tissue, and (ii) a thin film of water forms on the spine surface, where arriving droplets coalesce with the thin film, which drains down the spine surface under the influence of gravity. The pronounced straightening of the spines that we observe at early times suggests that imbibition initially dominates, whereas the slowing of spine uncurling, which occurs for *t*~100-200 s, indicates that the spines are almost saturated. Consistent with this early imbibition-dominated regime, capillary-rise experiments on detached spines show a wetting-front that rises with *h*_*f*_ ∝*t*^*1/2*^, in line with Lucas–Washburn dynamics in porous media (Supplementary Text, *47*). During fog exposure, water enters the spine radially and induces swelling, which manifests macroscopically as spine straightening. Consequently, the normalized spine height *H/L* provides a geometric proxy for mass uptake. Assuming approximately constant density of the swollen tissue, volumetric expansion scales with mass uptake. Experimentally, we observed *H* (*t*)∼ *t*^1/2^, and deduced *m(t)*∼ *t*^1/2^, consistent with capillary-limited imbibition (Supplementary Text).

For later times (*t*>250 s), we anticipate that drainage dominates over imbibition, and this is supported by our observation that water collection in the glass capillary at the spine base lags both the onset of fogging and the initial uncurling process (Fig. S1A). To understand the parameters relevant for drainage in more detail, we analysed the geometry and uncurling of the spines. The spines are porous and tapered, with a cross-sectional area that changes shape along their length (Fig. S1B), from elliptical near the tip to semi-elliptical and crescent near the base. This results in a gradient of tissue distribution, with more tissue on the dorsal than ventral side towards the spine base. Since the spines tend to be curved inwards towards the apical centre on the plant, this exposes each spine from a different relative angle in a laminar stream of fog. We tested the effect of spine orientation on the water collection rates parallel to the direction of fog, comparing ventral and dorsal exposure of a single spine (Fig. S1C). Initially, at *t* < 300 s, the spine collects more water when fog hits the dorsal side, which has a larger surface area due to the transversely curved tissue. However, on timescales of *t* > 600 s, the collection rates converge, indicating similar collection efficiencies for dorsal and ventral exposure. In addition to Fig. 1D, these experiments show directly that straightening increases water collection rates. The effect is particularly apparent for ventral exposure; with mean collection rates increasing from 0.81 ± 0.29 mg/min for 0s < *t* < 120 s to 1.78 ± 0.45 mg/min for 300 s < *t* < 600 s (means ± SD with *n*=5 repetitions for each orientation; Fig. S1C) due to straightening of the spine in a constant stream of fog. A closer alignment with gravity increases the hydrostatic pressure gradient in the thin surface film and promotes drainage.

Regarding the timescales of drainage, we made the following observations: the spines initially absorb the fog water that collects on their surface and imbibition causes swelling and expansion of cell walls. Swelling causes spines to bend towards the dorsal side, uncurl and straighten up on a similar timescale to capillary imbibition (Fig. 1C, S2). By straightening, the thin film of water on the surface of the spines aligns increasingly with the direction of gravity (Fig. 1B, S2A), driving the flow down the surface of the spine. In experiments, the timescales of water collection (*t*~1800 s, Fig. 1D) are much greater than the timescales for drainage (*t*~250 s, Fig. S1A). So, we assume that steady-state drainage occurs, and the volume (*V)* of surface run-off is given by the product of flow rate (*Q*) and time (*t*), *V ~ Qt*. Since the flux of a thin film of water down a curved plane scales with the inclination angle (*α*) as *Q ~ sin(α)* (Supplementary Text; (*48*)) and the inclination angle is approximately *α ~ arcsin(H*_*1800*_*/L)*, where *L* is the arc length of a spine, we find that *V ~ H*_*1800*_*/L* for *t*=1800 s (Fig. 1D).

### Tissue properties for swelling and shape morphing

The spine tissue shows hygroscopic expansion, like a cellulose sponge (*47*). Generally, cactus spines consist of dead libriform fibres and a sclerified epidermis at maturity (*37, 49, 50*). In *T. alonsoi*, the spine epidermis shows large fissures, which are mainly oriented perpendicular to the longitudinal spine axis and interrupt the epidermis entirely (Fig. 2A-B, Movies S3-S6). In the dry state, these epidermal fissures have an equivalent pore radius *r*_*eq*_ = 14.7 μm and contribute to the surface porosity of 16.4 % (corresponding to Area_pore_ /Area_total_). This porous surface promotes capillary imbibition, which is required for hygroscopic expansion of the spine tissue. In the dry state, the epidermal cell walls are folded, and cells have a larger diameter than the fibres of the core (Fig. 2B). Raman imaging of the epidermal cell walls reveals clear bands of cellulose and pectin (with a low degree of esterification, Fig. S3); with a strong pectin enrichment in the outer periclinal wall of the epidermis (Fig. 2C). Generally, cellulose and pectin play a key role during epidermal wall formation and for inducing folds in cell walls (*51*). The tissue contains water-soluble flavonoids with a strong and characteristic Raman band near 1572 cm^-1^ (Fig. S3C). A cuticle covers the spine surface and consists of a mixture of fatty acid esters and aromatic compounds (Fig. 2C, S3C). The core tissue of the spines consists of longitudinally oriented fibres with distinct secondary cell wall thickenings rich in cellulose and flavonoids (Fig. 2D). Cross-sections of the fibres reveal slightly compressed and folded cell walls in the wet state (Fig. 2D).

**Fig. 2.**
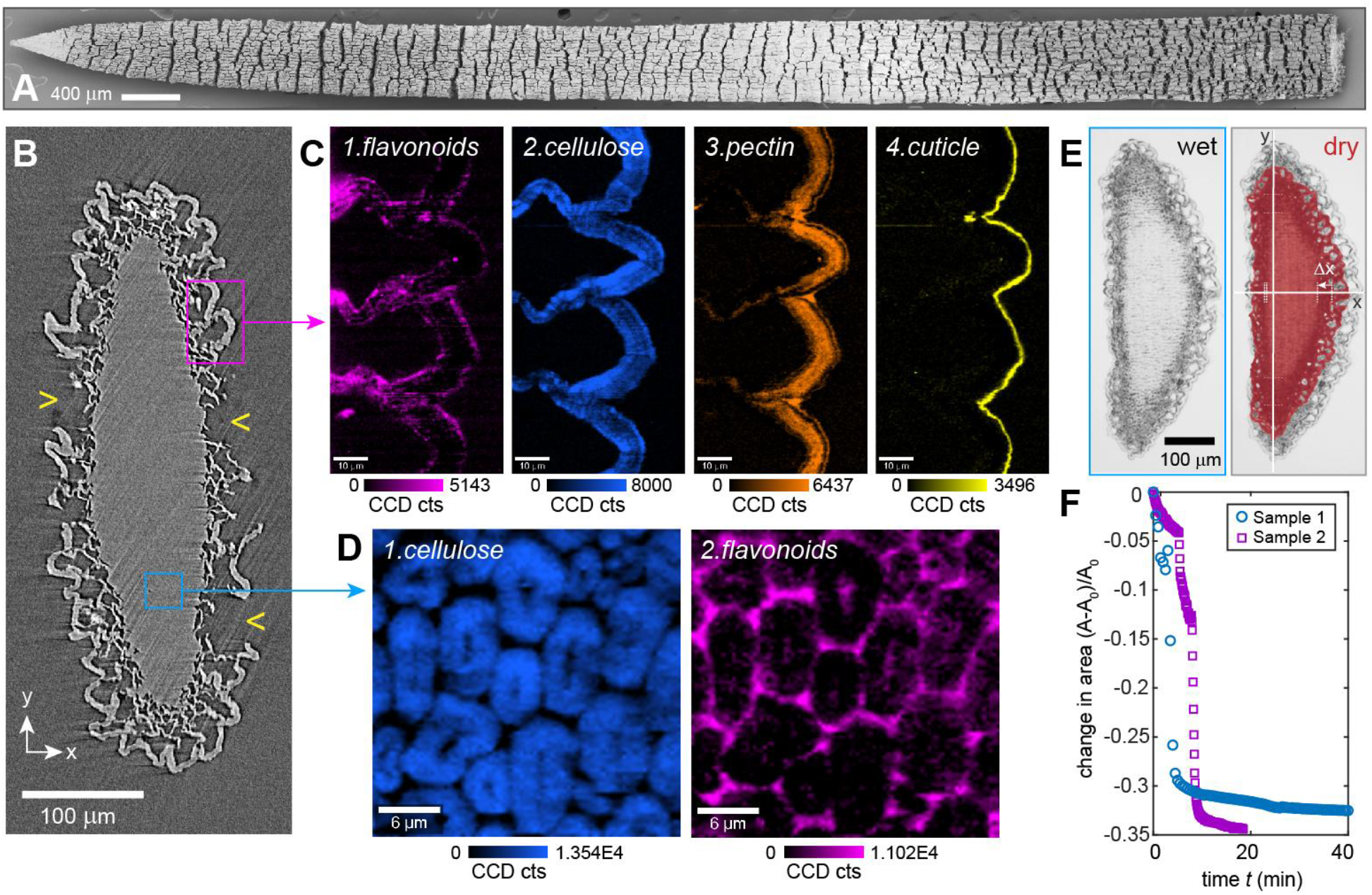
Microstructure, composition and shrinkage of the spine tissue. (**A**) SEM image of the spine surface (ventral side), showing many transversal fissures in the epidermis (see also Movies S3-S6); (**B**) μ-CT cross-section of a dry spine in the tip region, showing folded epidermal cell walls and deep fissures exposing the fibrous core tissue (indicated by arrowheads). (**C**) Confocal Raman images of cross-sections, showing the distribution of the most prevalent compounds in the epidermis and (**D**) the core tissue. (**E**) Light microscopy images of a cross-section in the wet and dry state (overlay coloured in red). (**F**) Relative change in tissue area as a function of time during drying of two sample cross-sections, recorded with frame rates of 30 s (sample 1) and 5 s (sample 2).

During drying of initially wet tissue sections, the overall cross-sectional area decreased by 30-35% (Fig. 2E-F). This strong shrinkage is predominantly caused by the radial shrinkage of the core fibres, but it also includes the folding of cell walls in the epidermis upon dehydration. In succulents, cell wall folding and unfolding are typical processes within the living tissue as a response to changes in tissue hydration and enable large volumetric changes (*52*). In *T. alonsoi*, the spine tissue is non-living at maturity and the only continuous structure in which stresses can accumulate and cause spine deformation during swelling and shrinkage is the dense core. This core shows highly anisotropic shrinkage: in the transversal plane, shrinkage is near uniform and amounts to 19-24 % in all four major directions (+x, -x and +y, -y), as measurements of a cross-section from the mid region indicate (Fig. 2E). In the longitudinal direction, however, shrinkage is considerably smaller – the length of a spine segment decreased by less than 3 % when dried from the wet state. From observations of anisotropic shrinkage under drying, we infer that swelling also occurs predominantly in the transversal plane. We attribute this macroscopic effect to the anisotropic swelling of the fibres, which exhibit greater transversal strain than longitudinal strain as they swell, as is typically the case for slender objects that deform in response to changes in internal pressurisation (*53*).

This anisotropic expansion, along with a pre-curved spine geometry, is sufficient to drive uncurling. We demonstrate this by means of Finite Element Method (FEM) simulations (Fig. 3), using a homogeneous, curved and tapered beam with a semi-elliptical (D-shaped, Fig. 3A) cross-section that is pinned at the base and subjected to orthotropic expansion with strain ε. Informed by our experimental observations of anisotropic strain, the expansion was mostly restricted to the transversal plane, with strains ranging from ε_x_ = ε_y_ = 0.05 to ε_x_ = ε_y_ = 0.25 (i.e. 5-25 %), whereas longitudinal expansion was fixed at ε_z_ = 0.025 (i.e. 2.5 %). The numerical simulations show that the spine starts to straighten into a more vertical position as soon as material expands in the transversal plane (Fig. 3B), characterized by an increase in the cross-sectional area (Fig. 3C) and spine height (Fig. 3D), comparable to the biological spine (Fig. 1-2). The finite element simulations confirm that transverse swelling strain drives spine straightening and quantitatively link volumetric expansion to the observed increase in normalized height *H/L*. This establishes a direct mechanical coupling between mass uptake and geometric reconfiguration, supporting our earlier interpretation that the experimentally observed scaling *H(t)*∼ *t*^1/2^ reflects capillary-limited imbibition within the internal porous network.

**Fig. 3.**
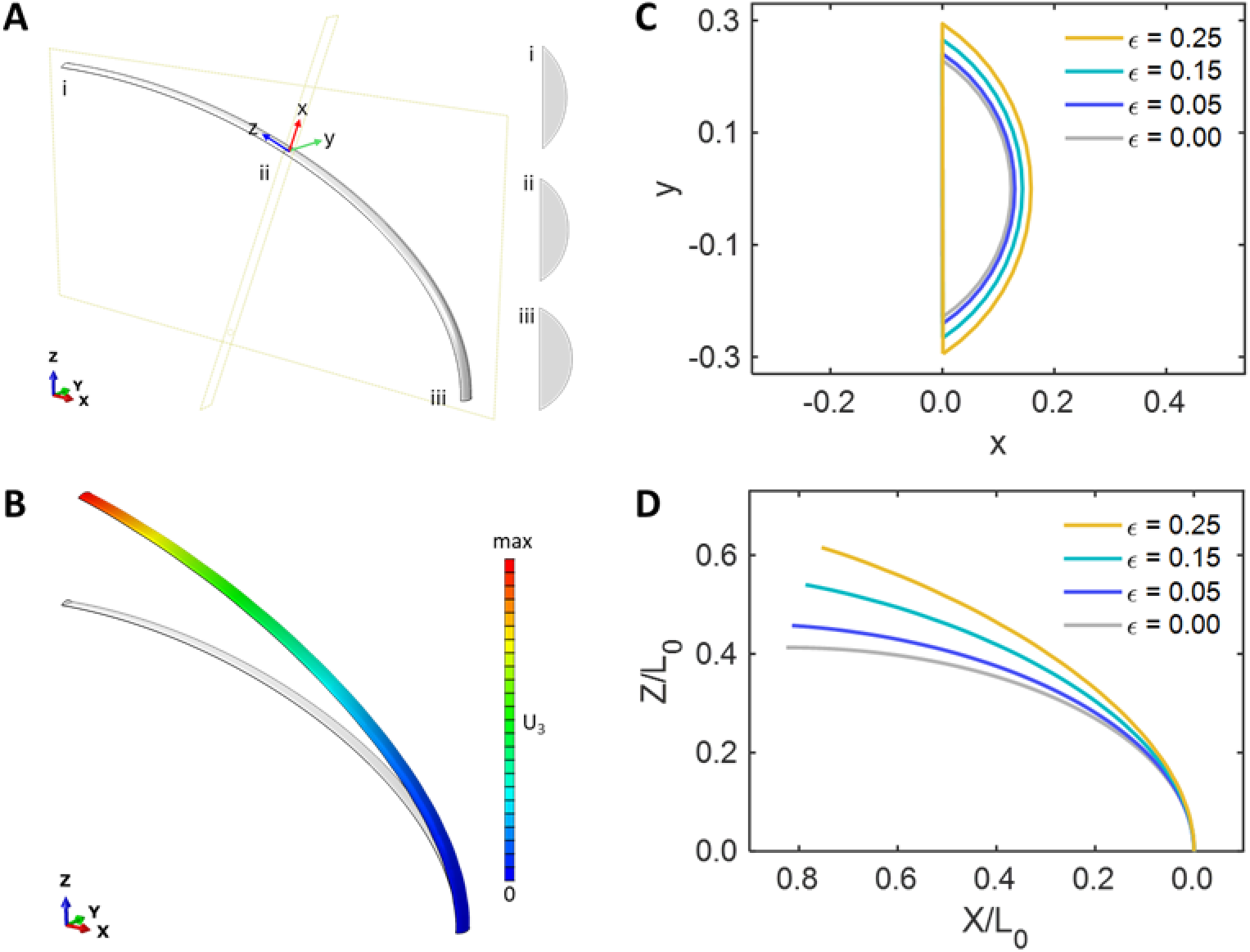
Simulating spine deformation via transversal expansion. (**A**) Spine geometry with cross-sectional shapes (i-iii) corresponding to the spine tip, mid and base, respectively in the global coordinate system (O_XYZ, shown at the bottom left) and the local coordinate system (O_xyz, shown at the middle of the spine) for the spatial expansion coefficients α_x_ = α_y_ = 10 α_z_. (**B**) Colour-coded displacement of the spine after expansion (with expansion strain *ε*_x_ = *ε*_y_ = 0.25 and *ε*_z_ = 0.025, see also Movie S7) and the original shape (grey). (**C**) Absolute geometry of the cross-section from the mid region (ii) for different transversal expansion strains (*ε = ε*_x_ = *ε*_y_) and *ε*_z_ = 0.025. (**D**) Longitudinal spine positions (lower side, absolute values) for different transversal expansion strains (*ε = ε*_x_ = *ε*_y_) and *ε*_z_ = 0.025. L_0_ corresponds to the total length of the initial, undeformed spine.

Uncurling also occurs in pre-curved beams with uniform cross-sectional geometries (Fig. S4), suggesting that the swelling ratio and initial longitudinal curvature have a greater influence on uncurling than the cross-sectional shape. Furthermore, our simulations demonstrate that dorsoventral expansion is the key driver behind uncurling as it generates a bending moment that can straighten the spine even in the absence of lateral expansion (Fig. S5). The straightening of the spine is therefore analogous to the tendency of a curved tube to straighten under internal pressurization (*54*) – a phenomenon that is commonly employed in Bourdon pressure gauges.

### Spine anchorage in a suberized tissue

Each spine grows from an axillary bud, known as an areole (*43*), and remains anchored after development (Fig. 4A). In this region, the tissue structure and chemical composition gradually change, as revealed by Raman imaging (Fig. 4B-D): at the spine base, the fibres in the core show thinner secondary cell walls (Fig. 4C, consisting mainly of cellulose and phenolics) than in the mid and tip regions of the spine and the primary walls are suberized (Fig. 4D). Furthermore, between the spine base and the living tissue of the areole lies a dense tissue termed ‘attachment layer’ (AL in Fig. 4B), consisting of compressed cells with thin walls that are strongly folded and suberized (Fig. 4E-G). The suberized tissue (AL) is directly connected to the living plant tissue via thick, pectin-rich cell walls that enclose many calcium oxalate crystals (Fig. 4F-G). The Raman spectra of the cell walls from the AL share all major bands with those of cell walls found in the bark of the cork oak *Quercus suber* (Fig. 4G), known to contain mainly suberin. Spectral reconstruction worked well with 5-hydroxyferulic acid and various fatty acids (e.g. Fig. S6), which is in accordance with the feruloylation reactions reported during suberization (*55*). The chemical and morphological features of this suberized tissue show many similarities with the (wound) periderm in other taxa, in which suberin lamellae are deposited outside of the primary cell wall (*56*). Generally, suberin has played an important role as water-proofing biopolymer during plant evolution (*57*), suggesting that the spine base and the attachment layer (Fig. 4B) are water impermeable.

**Fig. 4.**
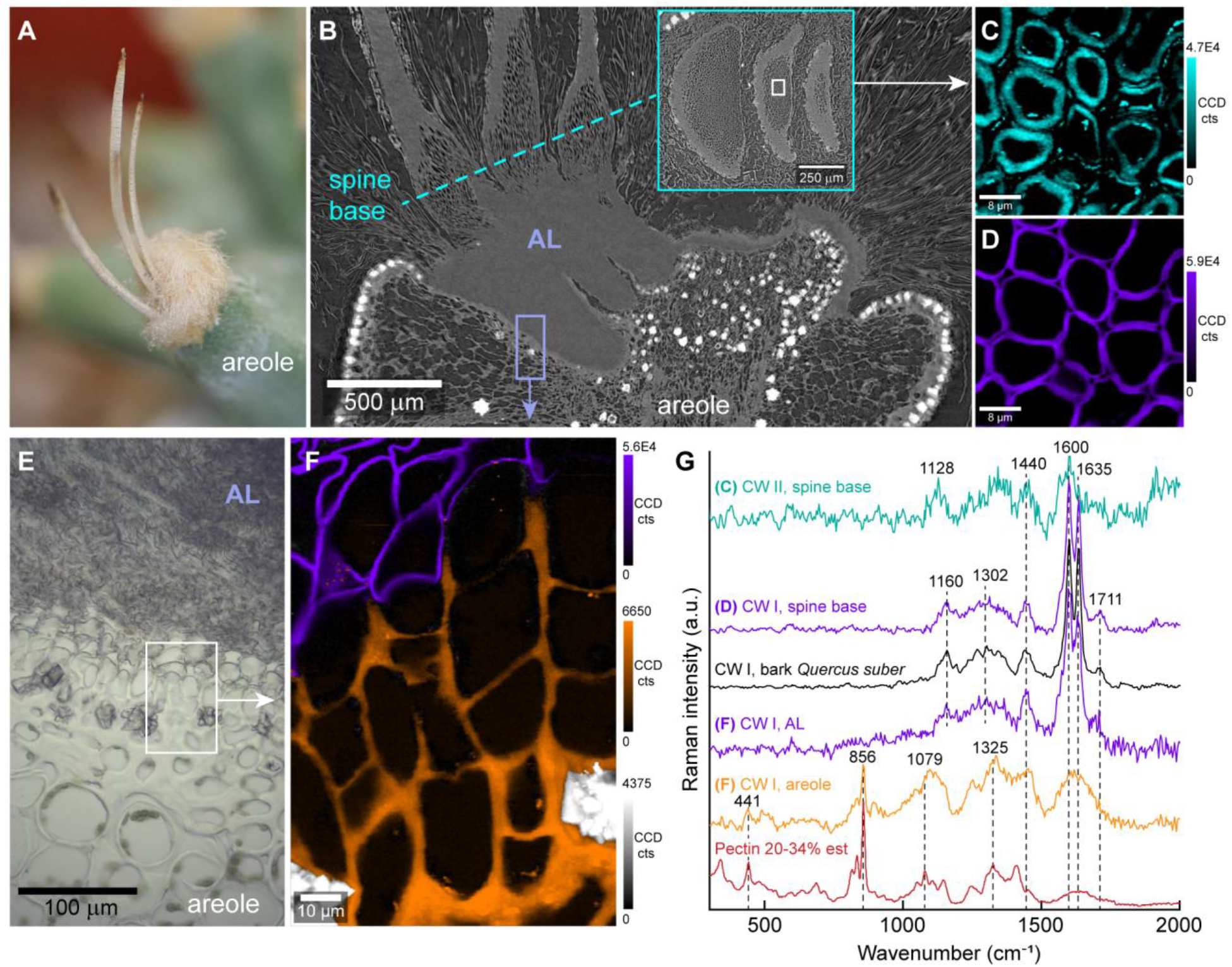
Spine anchorage and chemical composition of the attachment layer. (**A**) Photograph of an areole with three spines. (**B**) Longitudinal section (μ-CT image) of an areole with three spines (inset showing spine cross-sections at the base), revealing a dense attachment layer AL for spine anchorage. (**C-D**) Raman images of the core tissue, obtained from spine cross-sections at the base (box shown in inset of B), showing (**C**) pronounced secondary cell walls and (**D**) suberized cell walls. (**E**) Light microscopy image of a longitudinal section obtained from the transition zone between the attachment layer (AL) and the living tissue of the areole. Box illustrates an exemplary measurement area for (**F**) Raman imaging of the transition zone. The composite image includes suberin (purple), pectin-rich cell walls (orange) and calcium oxalate crystals (grey). (**G**) Raman spectra of the measurement areas and reference spectra from the bark of the cork oak and citrus pectin (20-34% esterified). CW: cell wall.

We verified this by stem water analysis from cactus plants that were exposed to deuterium-labelled fog, which should reveal a deuterium-enriched signal in waters extracted from cortex tissues if the attachment layer was permeable to water. However, the extracted stem water samples from *T. alonsoi* (and from *Copiapoa humilis* with a non-porous spine epidermis for comparison) show similar δ^2^H values as non-fogged plants (Fig. 5), even when doubling the exposure time to fog. This suggests that both the spine attachment layer at the areole and the entire cuticle surrounding the cortex are water impermeable. During plant transpiration, fractionation of water occurs naturally, leading to an enrichment of heavy isotopes in tissues (*58, 59*). This fractionation, i.e. the enrichment of deuterium, is also visible in the stem water of *T. alonsoi* and *C. humilis* by the offset from the global meteoric water line (GMWL, Fig. 5), representing the positive correlation of δ^2^H and δ^18^O in natural meteoric water samples.

**Fig. 5.**
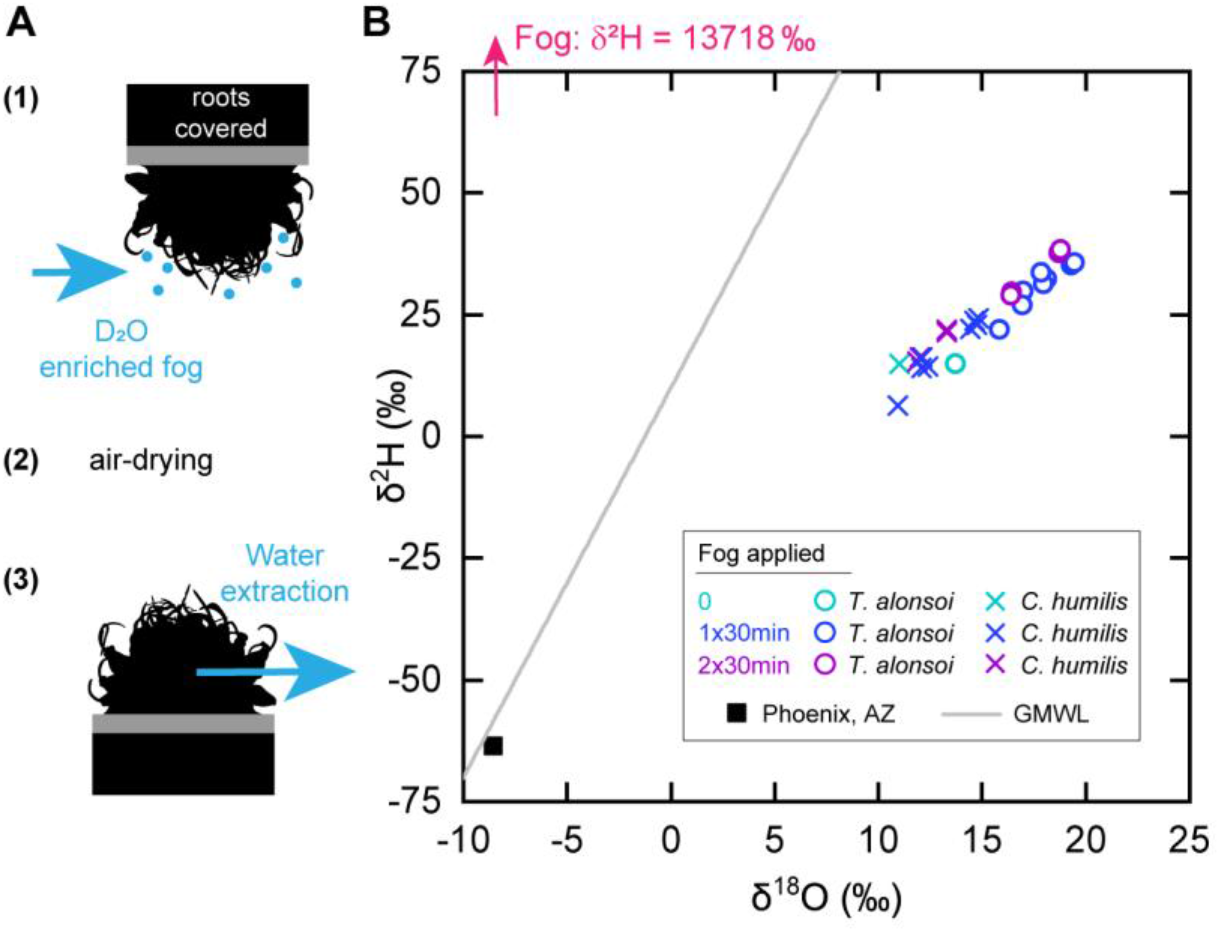
Stem water analysis of cactus plants exposed to D_2_O labelled fog. (**A**) Schematic overview of the fogging procedure with plants upside down and roots sealed. Water extraction was performed ∼1 week after fogging. (**B**) Stable isotope analysis of water samples by mass-spectrometry, revealing similar levels of deuterium enrichment in the cortex of *T. alonsoi* (and *C. humilis* for comparison), regardless of treatment (no fog, 1 × 30min and 2 × 30min). The global meteoric water line GMWL represents the positive correlation of δ^2^H and δ^18^O in meteoric water samples; a water sample from Phoenix, AZ is included and was used to prepare deuterium enriched water for fogging.

### The path of fog water

Based on our data, the path of fog water is as follows: (*i*) fog droplets hit the spine surface and are directly absorbed into the spine tissue via the fissures in the epidermis; (*ii*) water then further penetrates into the hygroscopic spine core by capillary forces, leading to cell wall swelling and spine straightening; (*iii*) excess water accumulates on the spine surface, forming a continuous liquid film that flows towards the spine base driven by gravity; (*iv*) and further propagates along the surface of the plant, also under the influence of gravity, (*v*) until reaching the soil and root system, where water can finally be absorbed into the living tissue.

## Discussion

In this study, we show that the porous and flexible spines of *T. alonsoi* undergo shape morphing when exposed to fog, characterized by uncurling and increased alignment with the direction of gravity, which leads to increased fog water collection rates. Fog harvesting may be crucial for survival of *T. alonsoi*, since fog forms regularly in the species’ habitat, contrasting the irregularly occurring rainfall and the limited soil water retention in the rocky substrate. Our work shows that hygro-morphing enables two distinct spine configurations: (i) under dry conditions, the spines are more curved and form a denser layer covering the apical meristem of the main shoot axis, which may protect it from excess sun light and mechanical damage (*60-62*), but also reduce drag and the risk of spine breakage (*63, 64*). (ii) When exposed to fog, the spines straighten into a more vertical orientation, which increases fog harvesting rates even at the relatively low wind speed of 0.2-0.3 m/s tested in our study. For lower wind speeds, other works have suggested that a more vertical orientation is also effective for intercepting droplets by diffusion, due to a higher exposure to vapour gradients (*65*). In an arid habitat, fog harvesting may confer a substantial advantage, particularly in the case of short-term advective fog harvesting.. For example, by considering a small change in spine height from H/L=0.6 to H/L=0.7 and 30 minutes of constant fogging, straightening can increase the amount of fog water collected up to 132% in a single spine (calculated based on the equation used in Fig. 1D). This suggests that the spines are adapted for fog and dew capture; but we found no evidence from isotopic labelling that water enters the inner cortex of the plant via the spines, as previously suggested (*30-33*). Instead, fog harvesting is restricted to the surface of the cortex in *T. alonsoi* and water absorption occurs only via the well-developed root system, which may also improve the uptake of soil nutrients. This contrasts with *Tillandsia sp*., in which fog water enters the cortex directly via the epidermal trichomes (*16*).

The Cactaceae family exhibits a large morphological and structural diversity of spines (*43, 66*). In the context of fog harvesting, at least two different types of spines have been described, including i) long, porous and upwardly oriented spines capable of hygro-morphing (this work), and ii) non-porous, stiff and conical spines with micro-structured surfaces (*30*). Our findings emphasize the role of structure and geometry for cactus spine functioning and raise the question of the evolutionary drivers of this diversity. The insights can serve as inspiration for shape-morphing materials and fog harvesting devices. Synthetic analogues of cactus spines have already been developed for fog harvesting (*21, 23-25*) and further advances may be achieved by using shape-morphing materials, porous layers for integrated water filtering, or distinct surface patterning to modulate droplet condensation, adhesion and transport (*26, 67-69*).

## Supporting information

Software code S1

Software code S2

Software code S3

## Acknowledgments

The authors thank Susanne Pamperl (University of Vienna), Martin Felhofer and Paraskevi Charalambous (BOKU), Dori Wolfe (UWYO), Noemí Hernández Castro and Ali Schuessler (DBG) for technical assistance, as well as Shahrouz Amini (MPI-CI), Peter Lechner and Lukas Schrangl (BOKU) for valuable discussions. We also thank Matthias Uhlig (Uhlig Kakteen, Germany) for supplying cactus plants. FB would like to acknowledge the Royal Society (URF\R1\211730) and ML would like to acknowledge the start-up funding from the University of Birmingham. JFL acknowledges support from NSF Grant POLS-2442203. This research was funded in whole or in part by the Austrian Science Fund (FWF) [grant DOI: 10.55776/ESP11]. For open access purposes, the author has applied a CC BY public copyright license to any author accepted manuscript version arising from this submission.

## Funding

Austrian Science Fund FWF (DOI: 10.55776/ESP11); JCH

Royal Society (URF\R1\211730); FB

National Science Foundation (POLS-2442203); JFL

## Author contributions

Conceptualization: JCH, FB, KRH

Methodology: all authors

Investigation: JCH, FB, QZ, MAG, TAO, JFL, DGW, ML, KRH

Visualization: JCH, FB, JFL, QZ, ML

Funding acquisition: JCH, NG

Project administration: JCH

Supervision: JCH

Writing – original draft: JCH, FB Writing – review & editing: all authors

## Data and materials availability

All data are available in the main text or the supplementary materials. Micro-CT scans can be downloaded from Zenodo (DOI: 10.5281/zenodo.14333372; Link for reviewers: https://zenodo.org/records/14333372?preview=1&token=eyJhbGciOiJIUzUxMiJ9.eyJpZCI6ImNiNzNlZjQ2LWZhNDQtNGY3Ni1hNWYxLTlhNjkwOGJiYzA3ZiIsImRhdGEiOnt9LCJyYW5kb20iOiJjNzQ2ZjZjMGFjZGJjMjlmMzdjNTYwNmRlZTg2NTlkZiJ9.i3cBgUKCg8wtHykopn4Dx2qm1D86EoY5-gPG9guJtbHJjnFTomj96FMo6p9VxK5dUPtx9qlEOcnHmJsjgKAO4g).

## Materials and Methods

### Plant material

For structural analysis *Turbinicarpus alonsoi* plants were purchased from Uhlig Kakteen (Germany) and kept indoors at the University of Natural Resources and Life Sciences Vienna. For stable isotope analysis, 6 individual plants of *Turbinicarpus alonsoi* and 6 individuals of *Copiapoa humilis* were purchased from Bach’s nursery (Tucson, Arizona, USA) and stored in a ventilated greenhouse at the Desert Botanical Garden in Phoenix (Arizona, USA) until use without any watering. Each plant had its own plastic pot and soil substrate.

### Videography of spine deformation and fog collection

All videos and photographs were taken with a macro-camera (EOS M10, Canon, objective EFS 35mm) inside a photo box (booth box 80 × 80 cm, Hakutatz) with maximum illumination. Typical experimental conditions were a temperature of *T*=23 °C and a relative humidity of *RH*=30-40 %. Whole plants and individual spines were fogged at a constant intensity with a terrarium fog generator (Repti Fogger, Zoo Med’s), using demineralized water. For the experiments with single spines, the attached bottle was dismounted and a volume *V*=200 mL was filled into the water reservoir, followed by closing the opening with parafilm to prevent evaporation. A weight of 800g was placed on top to stabilize the apparatus. The fog outlet was mounted 4.5 cm away from the central axis of the spine and the glass pipette for fog collection. The base of individual spines was glued onto a short piece of metal wire with as little superglue (Loctite, Henkel) as possible and placed into the narrower end of a Pasteur glass pipette (length *L*=230 mm, outer diameter *d*=1.43 mm; Roth, Germany) and mounted vertically with the thicker pipette end placed into a glass vial (volume *V*=10 ml, height *h*=45 mm; Roth, Germany) and held in place by stripes of a commercial cellulose sponge, allowing air to exit the pipette. To determine the amount of fog water that the spines collected, the weight of the spine (including the pipette and the glass vial) was determined in the dry state and subtracted from the weight after 30 minutes of fog exposure. Water droplets that accumulated on the outer surface of the glass pipette were carefully wiped away with a paper tissue before weighing. Each repetition of the experiment was done when the glass pipette and the spine were fully air-dried again and the dry weight identical to the previous repetition (rounded to the third decimal place of 1g). Samples were weighed with a microbalance (Sartorius R 160 P) and the weight recorded to four decimal places of 1g. To achieve different heights of the spine, the angle of the metal wire at the spine base was carefully adjusted with tweezers. The camera was placed at a 90° angle to the stream of fog and the spine. For scale, a background with pre-defined circle diameters was used. The same spine holder construction was used to measure the collection rates during ventral and dorsal exposure (with manual rotation of the spine) on top of a microbalance and at a distance of 4.5 cm between the central axis of the glass capillary and the fogger outlet. The same fog intensity was used as previously (corresponding to a velocity of roughly 0.2-0.3 m/s). The weight was recorded every 1 s with a TLC200 Pro time lapse camera (Brinno Inc.) placed in front of R 160 P microbalance (Sartorius).

### Video and photo analysis

The videos and photos of individual spines were processed in ImageJ (version 1.54, NIH), with videos first being converted from MP4 to AVI format. The spine height was measured manually with the rectangle tool and the length of spines measured with the freehand segmented line tool along the centre line of the spine.

Spine profiles were measured from calibrated images during fogging experiments using custom-built edge detection techniques in Matlab (MathWorks Inc., Software code S1 provided in the supplementary materials section). The spine curvature was estimated by measuring the radius of a circle fitted to detected profile, and assuming that the spine curvature is the reciprocal of the radius of curvature, *κ = 1/R*. Similarly, to determine the angle θ, we processed the images using a custom Python code (Software codes S2-S3 provided in the supplementary materials section) designed to detect the shape of the spines. For each image, we identified the edges of the spine and extracted its centreline. Along this centreline, we computed the local tangents and determined the angle θ between each tangent and the vertical axis (see Figure S2).

### Scanning electron microscopy

Straight spines from the apical region of the plant were harvested, air-dried and placed on top of a sample holder for scanning in an Apreo-SEM (FEI, ThermoFisher Scientific) with a voltage *U*=1.00 kV and a current *I*=0.10 nA. The entire spine was imaged with multiple overlapping images at the same magnification, which were then used to generate a stitched image afterwards using the automated photomerge command in Photoshop (Adobe).

### Micro-computed x-ray tomography

Individual spines, as well as entire areoles (cut with a scalpel and air-dried), were scanned with an EasyTom 150/160 system (RX solutions), using a Hamamatsu nano focus tube, set to a current of *I*=60A, a voltage *U*=170-200 kV and acquiring *n*=1440 projection images. Scans were reconstructed with the software Xact2 (RX solutions) and exported as z-stacks.

Overview scans were performed with a MicroXCT-200 (Zeiss Microscopy) with an exposure time *t*=10 s, a tube voltage *U*=35 kV, a current *I*=200 μA and a total of *n*=753 projection images per scan. Reconstruction of the 3D data was performed with the software XMReconstructor (version 8.1, Zeiss). All images were analysed in ImageJ (version 1.54, NIH).

### Confocal Raman imaging

Cross-sections of the spines were cut in a CM 3050 S cryo-microtome (Leica) with a thickness of 12 μm at a temperature of *T*=-12 °C. Sections were then transferred to a glass slide, immersed in H_2_O, covered with a high precision cover slip, and sealed with commercial nail polish. The tissue was measured with a confocal Raman microscope (Alpha300RA, WITec, Oxford Instruments), equipped with a 100× oil objective (NA=1.4, Zeiss Microscopy) and using a linear polarized Sapphire laser with a wavelength λ=532 nm (Coherent). Raman scattering was detected with a 50 μm optic multifibre connected to a spectrometer (UHTS-300 VIS, WITec, Oxford Instruments) using a grating of 600 g mm^-1^ and a CCD camera (DV401BV, Andor). Images were acquired at full laser power (P=44.8 mW) with an integration time of *t*=0.05s. Data analysis was performed with WITec Project 6 PLUS, using base-line corrected spectra in the spectral range from 220 cm^-1^ to 2000 cm^-1^. Individual sample components were identified by the integrated True Component Analysis (with negative weighing factors disabled and averaged spectra) in combination with WITec TrueMatch (version 6), using our own reference database and the Biochemicals database of ST Japan 5.2, containing 4795 reference spectra.

### Light microscopy

Cryo-cut spine cross-sections with a thickness of 12 μm were placed on top of a glass slide with a droplet of distilled water, covered with a glass cover slip and left to dry out on a light microscope (Labphot-2, Nikon) while images were automatically acquired every 30s or every 5s. The images were only used if the section deformed freely, without sticking to the glass, which was identified by visual inspection. Suitable images were then processed in Matlab (MathWorks Inc.) using custom-built edge detection techniques to determine the section boundary and to measure the cross-sectional area of the section for every image frame.

### Numerical simulations

We conducted finite element method (FEM) simulations using the commercial software ABAQUS to model expansion-induced deformation of the spine. The structure was discretized with 8-node linear brick elements with reduced integration and hourglass control (C3D8R). Mesh density was refined until the deformed shape and centreline displacement became mesh independent. The material was modelled as isotropic and linearly elastic, with *E* = 100 MPa and *ν* = 0.3 (since the deformation is governed primarily by geometry, these properties only scale the stresses generated by expansion and do not influence the deformation pattern). Volumetric expansion was implemented through the thermal-expansion feature with *orthotropic* coefficients aligned to local material orientation. The *x, y*, and *z* directions were defined to coincide with the thickness, width, and arc (length) directions of the beam, respectively. A unit temperature change (Δ*T*) was prescribed, and the thermal-expansion coefficients *α*_x_, *α*_y_, *α*_z_ were assigned to match the target expansion *ε*_x_, *ε*_y_, *ε*_z_. No additional mechanical loads were applied. Boundary conditions were applied by fully constraining the base nodes to remove rigid-body motion, while all other surfaces were left traction-free. Simulations were performed in a *Static, General* step with geometric nonlinearity enabled (NLGEOM = ON). Geometric dimensions were set to match those of the experimental specimens.

### Water extraction and stable isotope analysis

To test whether the spines transport water into the cortex of plants, fogging was performed at maximum intensity of the device using a solution highly enriched in deuterium consisting of demineralized water (*V*=3.780 L) and added D_2_O (*V*=8 mL; 99.9 % atom. D; Sigma-Aldrich Inc.), resulting in a D_2_O concentration of *c*=0.21 % and δ^2^H value of 13718 ‰. To prevent water leakage into the soil during fogging, the soil surface of the potted plants was fully covered with several paper tea bags (T-sac tea filters) filled with dried silica gel (Wisesorbent technology) and covered with 2 layers of parafilm. The entire pot was then placed into a plastic bag and taped together around the plant cortex. Any potential entry points for water were sealed off with more parafilm and 2 cm-wide gardening plastic strips were taped on top of the pot to stabilize the construction. The plants were then turned upside down and placed into a holder in the fume hood for fogging with the deuterium-enriched fog water. To test a potential dose-dependency, individual plants were fogged (Repti Fogger, Zoo Med’s) for a duration of 1 × *t*=30 min and others for 2 × *t*=30 min, both at maximum fogging intensity. After fog exposure, the plants were air-dried for *t*=1-2 h while still being upside down, then turned around, unwrapped, and checked for any leakage on the tea bags, followed by moving the plants to the greenhouse, where they were stored for about 1 week without any irrigation. Tissue extraction was then performed by puncturing the plant cortex horizontally with a sharp metal cylinder (diameter of *d*=0.45 cm), extracting two tissue samples from each plant with *m*=0.50-0.75 g for each sample, which was transferred into a plastic vial and stored in the freezer. The tissue of non-treated plants was already extracted a week prior (before any fogging) and stored in plastic vials in the freezer. All tissue samples were transported to the stable isotope facility of the University of Wyoming for cryogenic water extraction and mass spectrometry. Tissue water was extracted on the cryogenic vacuum water extraction line by transferring the tissue samples into N51A glass vials (ThermoFisher Scientific), which were then placed into larger sample glass vials (PYREX 25×200mm, Corning Inc.), covered by glass wool and attached to the vacuum extraction line. As soon as the vacuum was stable, the sample vials were heated to T ≈ 100 °C. The evaporated water was then trapped in the collection glass tube by cooling with liquid nitrogen. Finally, the glass tube containing the extracted water was flame-sealed. The samples were stored in a refrigerator until further processed. The isotope-ratio mass spectrometer (Delta Plus IRMS, ThermoFisher Scientific) was connected to a high-temperature conversion elemental analyser via an open split interface (Conflo IV) and operated in continuous flow mode. The extracted water samples were loaded into vials with volumes of *V*=200 μl to *V*=2.5 ml, sealed with PTFE/silicone/PTFE septa caps and processed with a liquid autosampler (Thermo AI 1310) attached to a flash HT analyser (ThermoFisher Scientific), injecting *V*=1 μl of sample water into the TC/EA column (filled with glassy carbon and held at a temperature of *T*=1420 °C). The gas chromatographic column within the elemental analyser was held at *T*=85 °C and separates the produced carbon monoxide and hydrogen gas via the open split interface, transferring the gases to the mass spectrometer. For each sample, three injections occurred, and the last two values were averaged. The isotopic composition of oxygen and hydrogen were given relative to Vienna Standard Mean Ocean Water (VSMOW) and SLAP scale, with 0 ‰ and -55.5 ‰ for oxygen and 0 ‰ and -428 ‰ for hydrogen, respectively. Normalization was performed with internal, quality-controlled reference materials.

The global meteoric water line (GMWL) was defined as follows:

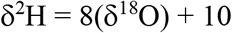

