## Supplementary Materials for "Humidity-driven shape morphing enhances fog harvesting in porous cactus spines"

**Content**

**This pdf file includes:**

Supplementary Text
SI References
Figures S1 to S6
Movie S1 caption
Movie S2 caption
Movie S3 caption
Movie S4 caption
Movie S5 caption
Movie S6 caption
Movie S7 caption

**Other supporting materials for this manuscript include the following:**

Movies S1 to S7
Software codes S1 to S3

### Supplementary Text

1. Properties of the fog:

The generated fog had a velocity of $v$=0.2-0.3 m/s, as calculated from video image analysis. The droplet diameter covered a range from $d$=10-60 µm, with a median diameter $d_{median}$=23 µm based on $n$=107 droplets imaged on a glass slide via a stereo microscope and analysed in ImageJ using thresholding and the particle analysis tool.

2. Physical description of imbibition and swelling:

a. *Capillary imbibition dynamics:* We conducted time-resolved capillary-rise experiments by dipping the tip of a spine into liquid water and tracking the height of the wetting front $h_f(t)$ as a function of time. Typically, this capillary rise height $h_f(t)$ in a porous medium follows a square root temporal scaling, $h_f(t) = a\,t^{1/2}$, with $a = 1.24\,\text{mm/s}^{1/2}$(Fig. R1). For early-stage imbibition in porous media, Lucas-Washburn scaling [1] predicts $h_f(t) \sim \left(\frac{\sigma R_{\text{eff}}}{\mu}\right)^{1/2} t^{1/2}$, where σ = 72 mN/m is the surface tension of water, μ=$10^{-3}$ Pa*s is its viscosity, and $R_{\text{eff}}$ is a hydraulic effective length scale governing the viscous–capillary. Combined with experimental data, this yields a calculated effective pore radius of $R_{\text{eff}} \sim 21$ nm.

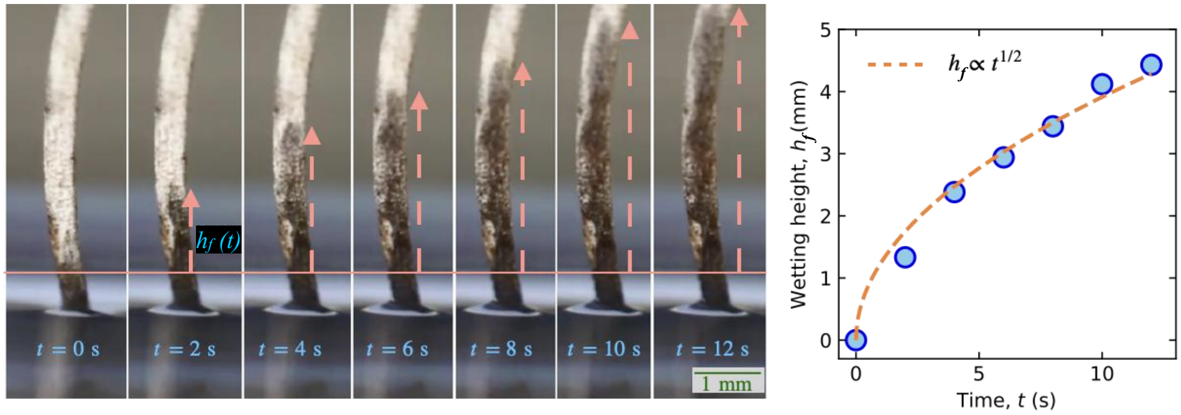


**Fig. R1: Capillary imbibition dynamics in a porous cactus spine of *T. alonsoi*.** *Left*: Time-lapse images showing spontaneous capillary rise of water along the spine after initial contact with the liquid reservoir ($t$ = 0–12 s). *Right*: The measured wetting height follows a square-root scaling $h_f \propto t^{1/2}$, consistent with Lucas-Washburn capillary imbibition in porous media.

b. *Internal structure and transport heterogeneity:* Cross-sectional imaging reveals a hierarchical internal architecture (Fig. 2, main text), characterized by larger outer voids and a denser inner matrix. Precise quantification of internal pore dimensions was not feasible. However, the observed structural heterogeneity indicates that transport occurs through a connected multi-scale porous network rather than through uniform cylindrical pores. In such heterogeneous systems, the parameter $R_{\text{eff}}$ extracted from the previous section (Washburn scaling) does not correspond to a geometric pore diameter. Instead, it reflects the combined influence of capillary pressure, permeability, tortuosity, and constrictions within the flow network. As shown by the hierarchical internal architecture of the spine (Fig. 2, main text), transport occurs through a multiscale structure in which flow resistance is governed by the most restrictive internal pathways rather than by the larger surface voids. The extracted $R_{eff}$ should therefore be interpreted as a lumped hydraulic scale characterizing the dominant flow resistance of the network.

c. *Directional imbibition and coupling with swelling:* During fog harvesting, water imbibition occurs radially and saturation of the spines occurs at $t \leq 200$ s (Fig. 1, main text). We used results from FEM simulations (Fig. 3, main text), which show that the transverse linear swelling strain $\varepsilon$ is directly related to the height change of the spine $H(t)$. Assuming constant density of the swollen tissue, $\frac{m}{m_0} = \frac{V}{V_0}$, where $V_o$ is the volume of dry spine at t=0 and $V$ is the volume of spine after swelling at time *t*. For orthotropic expansion with dominant transverse strain $\varepsilon_x = \varepsilon_y = \varepsilon$ and smaller longitudinal strain $\varepsilon_z = 0.025$, (Fig. 3, main text) the volumetric expansion is $\frac{V}{V_o} = (1+\varepsilon_x)(1+\varepsilon_y)(1+\varepsilon_z) = (1+\varepsilon)^2(1+0.025) = \frac{m}{m_o}$. Since FEM simulations show that transverse strain $\varepsilon$ drives straightening and increases the normalized spine height $H/L$, we infer a geometric coupling between swelling and height gain: $\frac{v}{v_o} = \frac{m}{m_0} = 1.025(1+\varepsilon)^2 \Rightarrow \frac{m}{m_0} \sim f\left(\frac{H}{L}\right)$. In Fig. R2, we plotted $\frac{m}{m_0}$ against $\frac{H}{L}$, showing a linear relationship between these two quantities.

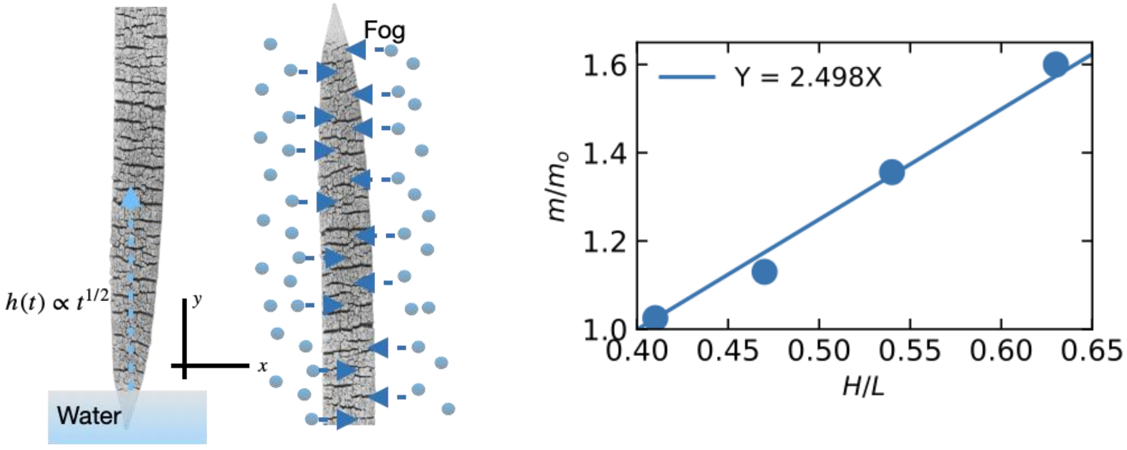


**Fig. R2: Directional imbibition in spines.** *Left*: Schematic illustrations of vertical capillary rise under reservoir contact and radial imbibition of water during fog exposure. *Right*: Normalized mass plotted against normalized height during fogging, and a linear fit.

d. *Modeling of lateral imbibition during fog exposure*: From our measurements, we observe that $(H(t) - H_o) \sim t^{\frac{1}{2}} \Rightarrow H(t) \sim t^{\frac{1}{2}}$ during the imbibition regime. Since we have established that $m(t) \sim H(t)$, it follows that $m(t) \sim t^{1/2}$ as well, as observed in Fig. R3. By analogy with vertical capillary-rise experiments, the lateral imbibition mass during early fog experiments can be described as $m(t) \sim t^{1/2}$.

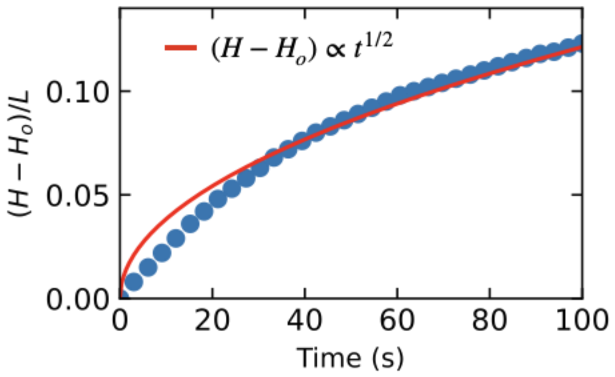


**Fig. R3:** Change in normalized spine height $(H - H_o)/L$ during early fog imbibition, showing $t^{1/2}$ scaling consistent with capillary-driven transport through the transverse pore network.

3. Analytical description of the flow along the outer surface of a curved thin plate:

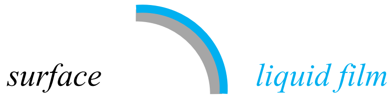


The flow of a liquid film along a curved surface at steady state can be described as follows; with

$Q$ = net volume flux of fluid down the surface
$x$ = flow direction, $z$ = axis always perpendicular to the flow direction,
$\alpha$ = inclination of the surface relative to the horizontal axis,
$v$ = fluid velocity,
$h$ = film thickness,
$\gamma$ = surface tension of the fluid,
$g$ = gravitational acceleration,
$\rho$ = density of the fluid,
$\mu$ = dynamic viscosity of the fluid.
p = pressure in the film
$\kappa$ = curvature of the surface

The boundary conditions are defined as $v = 0$ at the surface, $\frac{\partial v}{\partial z} = 0$ at $z = h$.
Neglecting velocity gradients in the flow direction and using the Navier Stokes equation yields

$$\mu \frac{\partial^2 v}{\partial z^2} = \frac{\partial p}{\partial x} + \rho g \sin \alpha$$

$$\rightarrow v = \frac{1}{2\mu}\left(\rho\, g \sin \alpha - \gamma \frac{\partial \kappa}{\partial x}\right) z^2 + c_1 z + c_2$$

with the boundary conditions $v = 0$ at $z = 0$; $c_2 = 0$ and $\frac{\partial v}{\partial z} = 0$ at $z = h$; $c_1 = h \frac{\gamma}{\mu} \frac{\partial \kappa}{\partial x} - \frac{h\, \rho\, g \sin \alpha}{\mu}$, and gives the volume flux

$$Q = \int_0^h v\, dz = \frac{\rho\, g \sin \alpha}{\mu} \frac{h^3}{3}, \text{ if } \kappa = const.$$

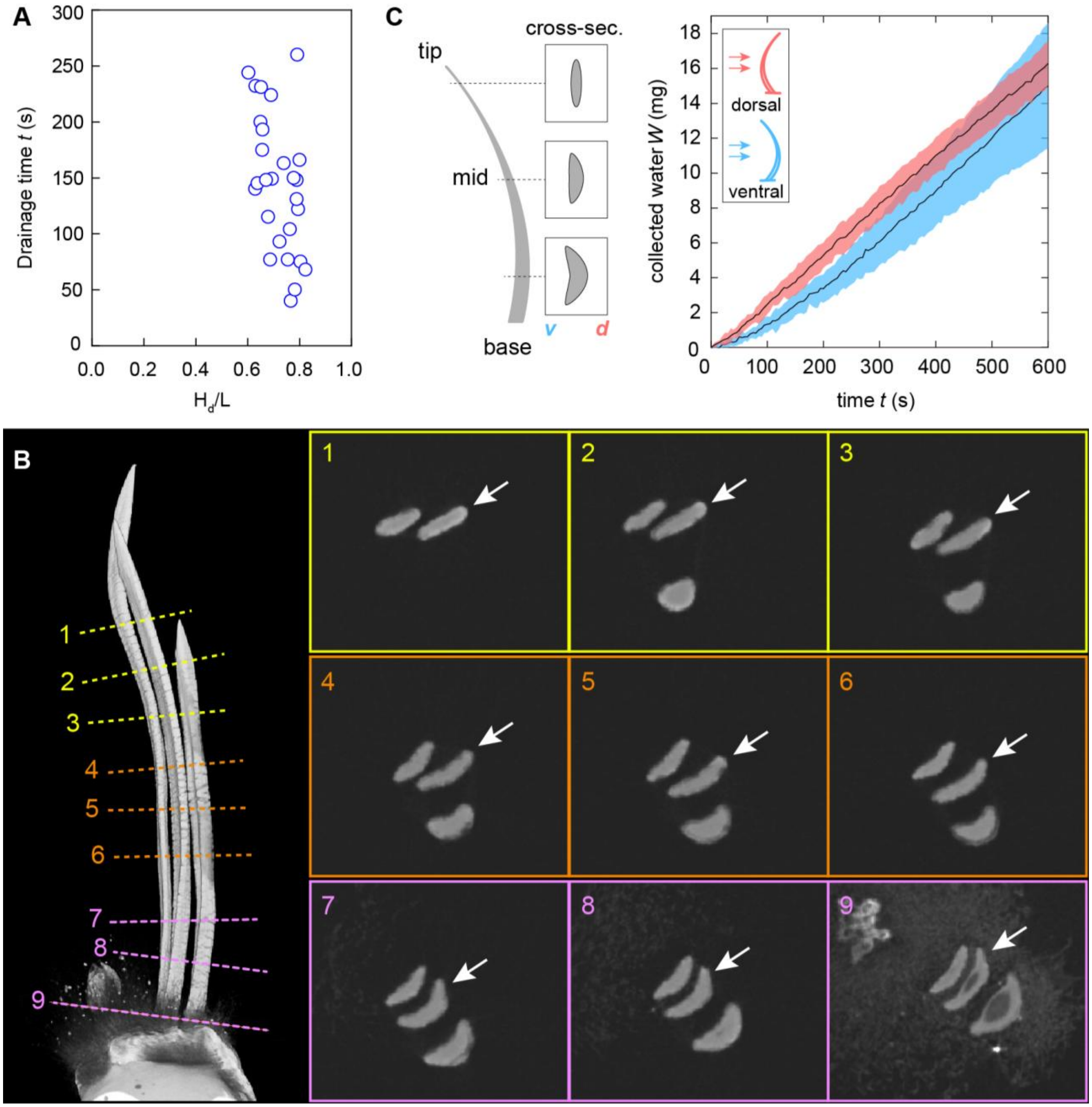


**Fig. S1.**

**Drainage time and effects of orientation on fog collection.** (**A**) Drainage time as a function of drainage height $H_d/L$ for ventral exposure of spines to fog (with a constant intensity). Drainage times were determined visually from video images, corresponding to the onset of water accumulation in the glass capillary. The drainage height $H_d$ is the height of the spine at the onset of water accumulation in the glass capillary. (**B**) Schematic illustration of spine geometry, showing exemplary cross-sections. (*v*: ventral; *d*: dorsal). (**C**) Collected water by spine 2 during fog exposure with its dorsal and ventral side (means ± SD; n=5 repetitions for each orientation). (**D**) Volume rendering of a micro-computed x-ray tomography scan of three spines attached to the areole. Lines indicate the position of each cross-sectional plane; the spine in the middle is marked by an arrow.

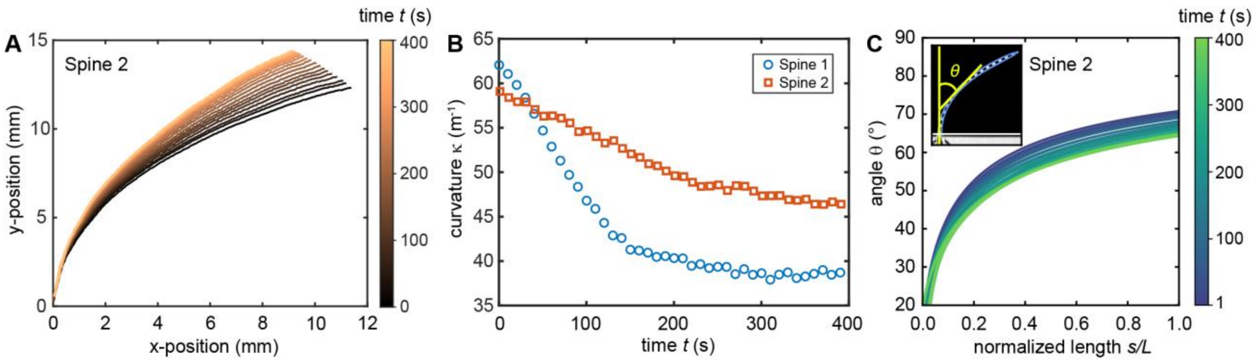


**Fig. S2.**

**Quantification of spine deformation.** (**A**) Absolute positions of spine 2 (mirrored) during straightening, characterized by a decrease in global curvature. (**B**) Global curvature $\kappa$ along the whole spine during fog exposure, which can be estimated by fitting a circle to the detected spine profiles and determining the reciprocal of the measured radius of curvature. (**C**) The angle $\theta$ is defined as the local deviation from the vertical axis at a given point $s$ along the centreline of the spine, described by the normalized length $s/L$ and as a function of time. The elliptical region near the spine tip is initially straight and remains straight during swelling. The straight tip region thus serves to amplify geometrically the height gain from uncurling of the mid region.

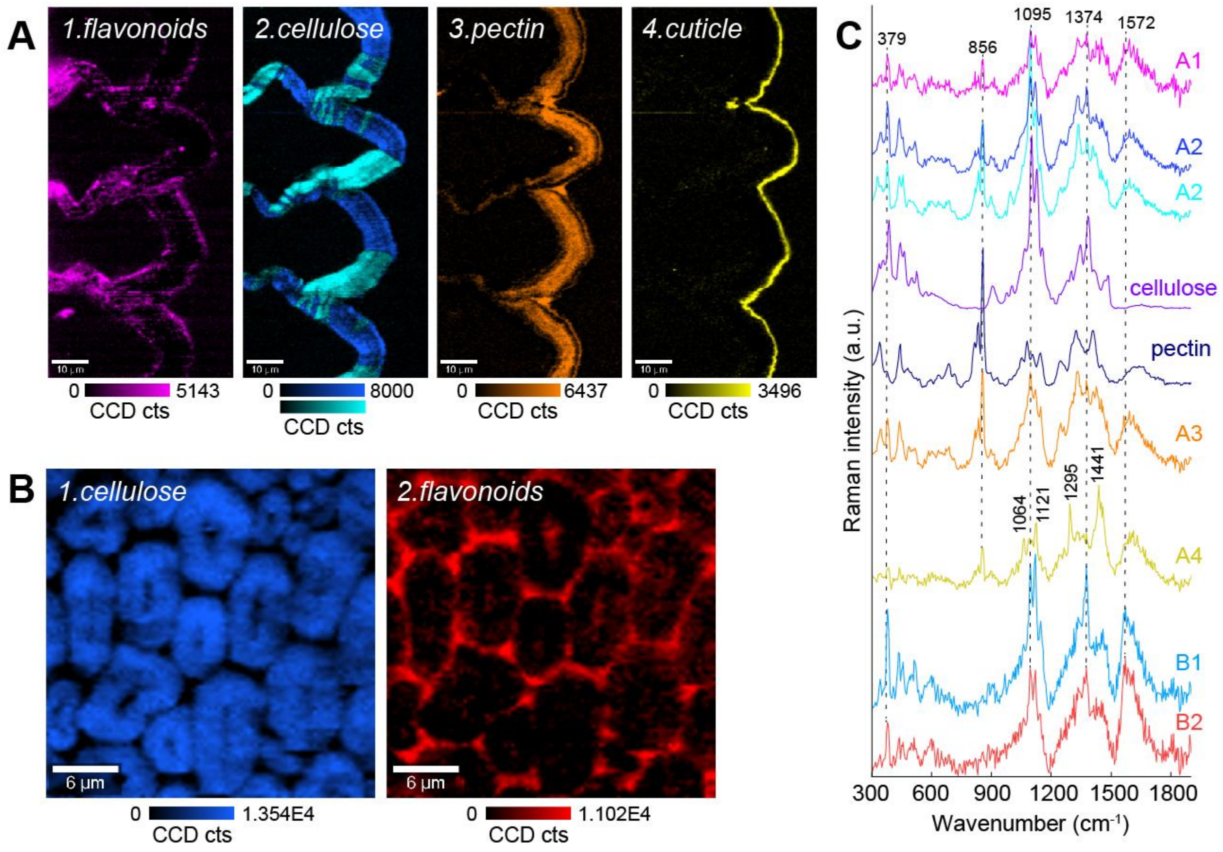


**Fig. S3.**

**Raman images and spectra obtained via True Component Analysis from cross-sections of the spine epidermis and spine core shown in the main text Fig. 2C-D.** (**A**) Intensity of individual components identified in the spine epidermis. The image of cellulose is a composite image based on two separate spectra, which were identified by the algorithm due to the changes in relative band intensity of cellulose as the orientation changes with respect to the laser polarization. (**B**) Intensity of individual compounds in the fibrous spine core. (**C**) Corresponding Raman spectra of the identified components in A and B via True Component Analysis, as well as reference spectra of cellulose and citrus pectin (20-34% esterified) for comparison. Characteristic band positions are indicated.

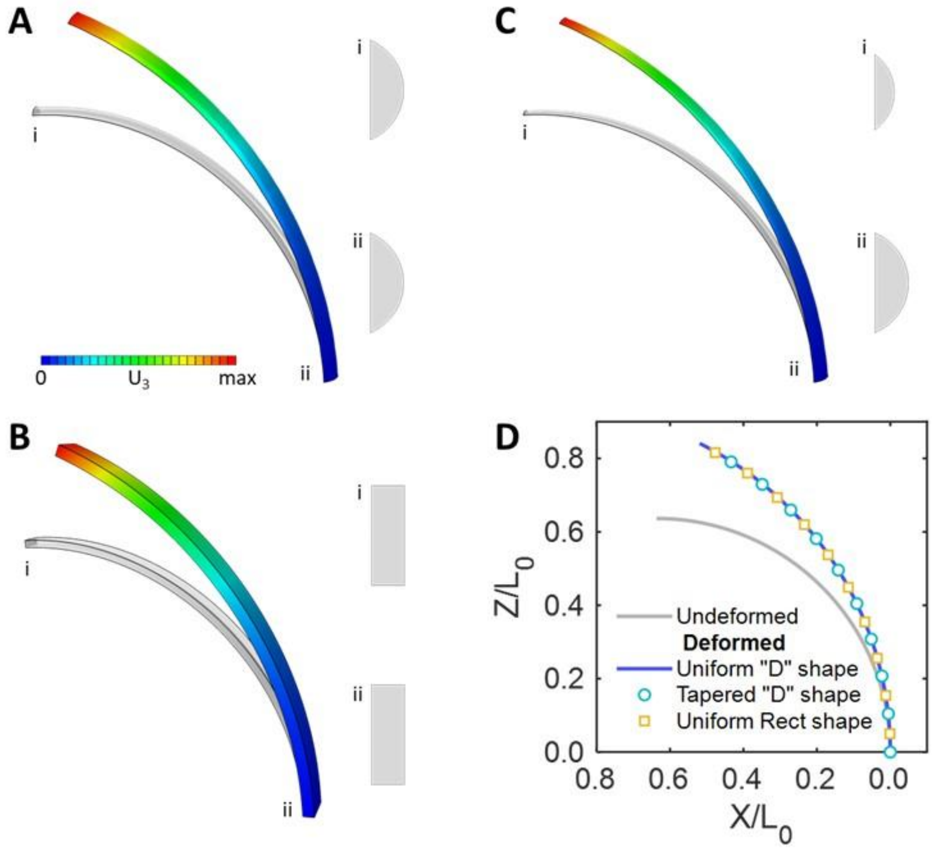


**Fig. S4.**

**Effect of cross-sectional shape on the displacement of a pre-curved beam under identical imposed expansion** (with strains $\varepsilon_x = \varepsilon_y = 0.40$, $\varepsilon_z = 0.04$). Original geometry is shown in grey; deformed configuration in colour (colormap indicating the displacement in Z direction). (**A**) Uniform "D"-shape cross-section. (**B**) Uniform rectangular cross-section. (**C**) Tapered "D"-shape cross-section (tip smaller and thinner than the base). (**D**) Normalized centreline profiles for the three designs in (A–C), showing nearly identical displacements under the same loading. $L_0$ corresponds to the original length of the spine in the undeformed state.

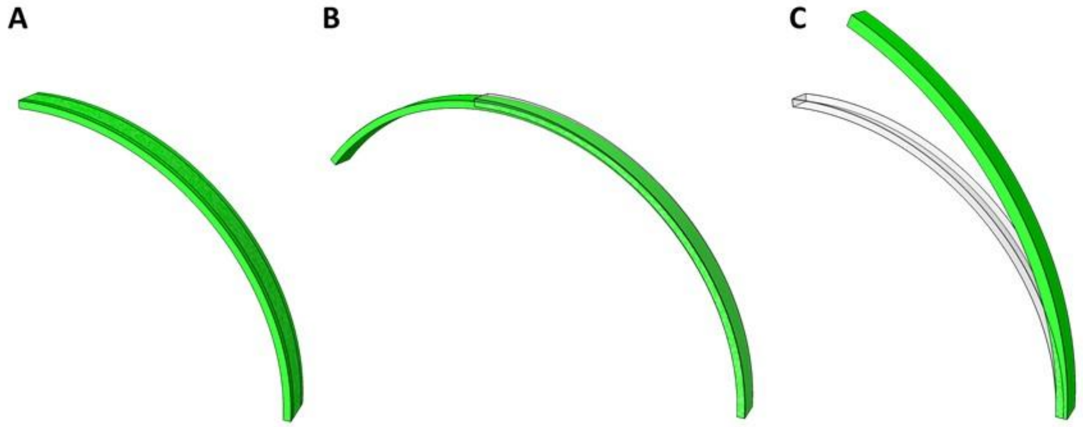


**Fig. S5.**

**Effect of the direction of expansion on the deformation of a pre-curved beam.** The undeformed state is shown in grey, and the deformed state in green. (**A**) For expansion strains $\varepsilon_y = 0.40$ and $\varepsilon_x = \varepsilon_z = 0.00$ the beam expands laterally, leading to a change in width. (**B**) For $\varepsilon_z = 0.40$ and $\varepsilon_x = \varepsilon_y = 0.00$ the beam elongates along the arc direction. (**C**) For $\varepsilon_x = 0.40$ and $\varepsilon_y = \varepsilon_z = 0.00$ the beam undergoes uncurling of the initial curvature, which is similar to the behaviour observed in the plant.

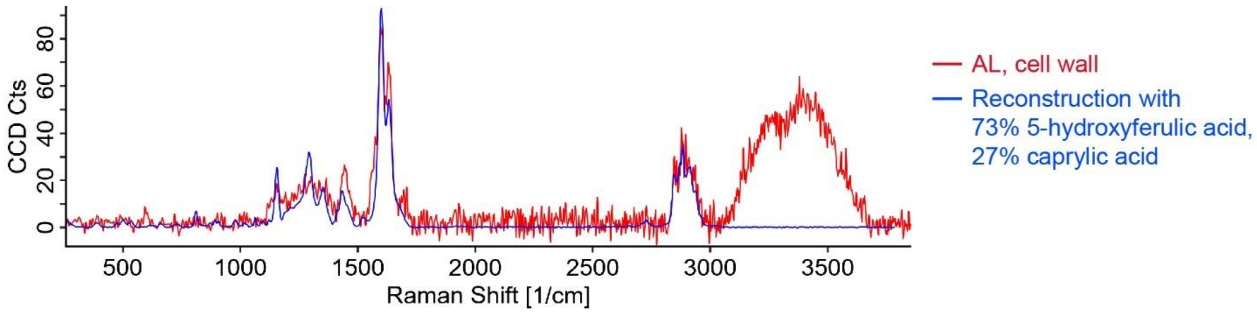


**Fig. S6.**

**Spectral reconstruction of the Raman spectrum identified in the suberized cell walls of the attachment layer (AL) of *T. alonsoi* (Fig. 4F-G, main text)** with 5-hydroxyferulic acid and caprylic acid. The ester band near 1711 cm$^{-1}$ points to an esterification of hydroxyferulic acid and fatty acids, which the model failed to reconstruct (possibly due to noise and a lack of suitable reference spectra). The bands from 3000-3600 cm$^{-1}$ arise from water ($H_2O$).

**Movie S1.**

*T. alonsoi* plant exposed to a stream of fog.

**Movie S2.**

Single spine (spine 1) of *T. alonsoi* exposed to a stream of fog. The diameter of the innermost circle corresponds to 3 cm. Time shown as min:sec.

**Movie S3.**

3D reconstruction of a *T. alonsoi* spine in the tip region (segment 1) based on high-resolution x-ray computed micro-tomography, showing the epidermal fissures on the ventral and dorsal side.

**Movie S4.**

3D reconstruction of a *T. alonsoi* spine in the mid region (segment 2) based on high-resolution x-ray computed micro-tomography, showing the epidermal fissures on the ventral and dorsal side.

**Movie S5.**

3D reconstruction of a *T. alonsoi* spine in the mid region (segment 3) based on high-resolution x-ray computed micro-tomography, showing the epidermal fissures on the ventral and dorsal side.

**Movie S6.**

3D reconstruction of a *T. alonsoi* spine in the base region (segment 4) based on high-resolution x-ray computed micro-tomography, showing the epidermal fissures on the ventral and dorsal side.

**Movie S7.**

FEM simulation of spine deformation with longitudinal expansion strain $\varepsilon_z = 0.025$ and transversal expansion strain $\varepsilon_x = \varepsilon_y = 0.25$, similar to the experimental values obtained from tissue shrinkage (Fig. 2E-F, main text).
