## Supplementary material for "Humidity-driven shape morphing enhances fog harvesting in porous cactus spines": Software code S2

# Code for spine centerline detection

import cv2
import numpy as np
import matplotlib.pyplot as plt
from scipy.interpolate import UnivariateSpline
from matplotlib import patches  # Import patches for drawing arcs

# Canny Algorithm for edge detection and finding topmost and bottom edge points
def calculate_edges(image):
    edges = cv2.Canny(image, 200, 500)  # Lower and upper thresholds for the hysteresis procedure in edge detection
    top_edge_points = {}                # Dictionary to store x-coordinate for each Y-coordinate for top edge
    bottom_edge_points = {}             # Dictionary to store x-coordinate for each Y-coordinate for bottom edge

    # Scan from top to bottom for top edge
    for x in range(edges.shape[1]):         # Loop iterates over each column (x) of the image
        for y in range(edges.shape[0]):     # Loop scans from the top (y = 0) to the bottom of the column
            if edges[y, x] > 0:             # Any pixel value > 0 recorded as edge points
                top_edge_points[y] = x      # This edge points are recorded in X for corresponding Y 
                break                       # After scanning is done break the loop
    
    for y in range(edges.shape[0]):  # Loop iterates over each row (y) of the image
        for x in range(edges.shape[1]):  # Loop scans from the left (x = 0) to the right of the row
            if edges[y, x] > 0:  # Any pixel value > 0 recorded as edge points
                bottom_edge_points[y] = x  # This edge points are recorded in X for corresponding Y
                break  # After scanning is done break the loop
                
    # Return 3 outputs: edges binary image, and 2 dictionaries that store edge points
    return edges, top_edge_points, bottom_edge_points   

# Draw top and bottom edge lines
def draw_edge_lines(ax, top_edge_points, bottom_edge_points):
    top_x_coords = [top_edge_points[y] for y in sorted(top_edge_points.keys())]
    top_y_coords = sorted(top_edge_points.keys())
    
    bottom_x_coords = [bottom_edge_points[y] for y in sorted(bottom_edge_points.keys())]
    bottom_y_coords = sorted(bottom_edge_points.keys())

    ax.plot(top_x_coords, top_y_coords, color='cornflowerblue', linewidth=0.7)  # Red color for the top edge line
    ax.plot(bottom_x_coords, bottom_y_coords, color='cornflowerblue', linewidth=0.7)  # Blue color for the bottom edge line

# Function to calculate the centerline between top and bottom edges
def calculate_centerline(top_edge_points, bottom_edge_points):
    centerline_points = {}
    for y in top_edge_points.keys():
        if y in bottom_edge_points:
            x1 = top_edge_points[y]
            x2 = bottom_edge_points[y]
            x = (x1 + x2) / 2  # Use '/' for floating-point division
            centerline_points[y] = (x, y)
    return centerline_points

# Smoothing centerline function
def smooth_centerline(centerline_points, smoothing_factor=3): 
    if not centerline_points:
        return []
    y_coords = sorted(centerline_points.keys())
    x_coords = [centerline_points[y][0] for y in y_coords]
    spl = UnivariateSpline(y_coords, x_coords, s=smoothing_factor)
    ys = np.linspace(min(y_coords), max(y_coords), 1000)
    xs = spl(ys)
    smoothed_points = [(x, y) for x, y in zip(xs, ys)]
    return smoothed_points

def draw_tangents_only(ax, center_points, num_tangents=1, tangent_length=120, thickness=1):
    x_coords = [p[0] for p in center_points]
    y_coords = [p[1] for p in center_points]

    poly_coeffs = np.polyfit(x_coords, y_coords, deg=5)
    poly_func = np.poly1d(poly_coeffs)
    poly_derivative = np.polyder(poly_func)

    middle_x = (min(x_coords) + max(x_coords)) / 1.35
    tangent_points = np.linspace(min(x_coords), max(x_coords), num_tangents - 1)
    tangent_points = np.insert(tangent_points, len(tangent_points) // 2, middle_x)

    for x in tangent_points:
        y = poly_func(x)
        slope = poly_derivative(x)
        length = tangent_length
        dx = length / np.sqrt(1 + slope**2)
        dy = slope * dx

        pt1 = (x - dx, y - dy)
        pt2 = (x + dx, y + dy)

        ax.plot([pt1[0], pt2[0]], [pt1[1], pt2[1]], color='gray', linewidth=thickness)

def draw_vertical_line1(ax, x_position, y_start=100, y_end=200, color='black', linestyle='--', linewidth=1):
    ax.plot([x_position, x_position], [y_start, y_end], color=color, linestyle=linestyle, linewidth=linewidth)

def draw_vertical_line(ax, position, color='black', linestyle='--', linewidth=1):
    ax.axvline(x=position, color=color, linestyle=linestyle, linewidth=linewidth)

def draw_horizontal_line(ax, position, color='black', linestyle='--', linewidth=1):
    ax.axhline(y=position, color=color, linestyle=linestyle, linewidth=linewidth)

def draw_shortest_distance_lines(ax, top_edge_points, bottom_edge_points):
    shortest_paths = []
    for top_y, top_x in sorted(top_edge_points.items()):
        min_distance = float('inf')
        closest_bottom_point = None
        for bottom_y, bottom_x in bottom_edge_points.items():
            distance = np.sqrt((top_x - bottom_x) ** 2 + (top_y - bottom_y) ** 2)
            if distance < min_distance:
                min_distance = distance
                closest_bottom_point = (bottom_x, bottom_y)
        if closest_bottom_point:
            shortest_paths.append(((top_x, top_y), closest_bottom_point))
    return shortest_paths

def draw_center_points(ax, shortest_paths):
    center_points = []
    for (top_x, top_y), (bottom_x, bottom_y) in shortest_paths:
        center_x = (top_x + bottom_x) / 2
        center_y = (top_y + bottom_y) / 2
        center_points.append((center_x, center_y))
    return center_points

def draw_centerline_curve(ax, center_points):
    if not center_points:
        return
    center_x_coords = [p[0] for p in center_points]
    center_y_coords = [p[1] for p in center_points]
    
    spl = UnivariateSpline(center_x_coords, center_y_coords, s=0.1)
    xs = np.linspace(min(center_x_coords), max(center_x_coords), 1000)
    ys = spl(xs)
    
    ax.plot(xs, ys, color='black', linewidth=1, linestyle='--')
    

def draw_arc_and_theta(ax, arc_center, theta_position, radius=10, theta_offset=0, arc_color='gray', theta_color='gray', arc_rotation=180):
    angle = np.radians(theta_offset + theta_position)

    arc = patches.Arc(arc_center, 2*radius, 2*radius, angle=200, theta1=0, theta2=np.degrees(angle), color=arc_color, linewidth=1)
    ax.add_patch(arc)
    
    theta_x = arc_center[0] + radius * np.cos(angle / 1.3)
    theta_y = arc_center[1] + radius * np.sin(angle / 0.1)
    
    ax.text(theta_x, theta_y, r'$\theta$', fontsize=12, verticalalignment='bottom', horizontalalignment='right', color='gray')


# Example usage
# image_path = '/Users/ovee/Downloads/f1_cleanup.jpg'
# larger_image_path = '/Users/ovee/Desktop/f2.jpg'

image_path = '/Users/ovee/Desktop/Image/Final plots/f1_cleanup.jpg'
larger_image_path = '/Users/ovee/Desktop/Image/Final plots/f2.jpg'

image = cv2.imread(image_path, cv2.IMREAD_GRAYSCALE)
larger_image = cv2.imread(larger_image_path, cv2.IMREAD_GRAYSCALE)

edges, top_edge_points, bottom_edge_points = calculate_edges(image)
centerline_points = calculate_centerline(top_edge_points, bottom_edge_points)

# Ensure the larger image has the correct dimensions
main_img_h = 450
main_img_w = 406
larger_image_resized = cv2.resize(larger_image, (main_img_w, main_img_h))

# Create a new image with the dimensions of the larger image and copy the larger image to it
combined_image = larger_image_resized.copy()

# Place the output image on top of the combined image
combined_image[0:image.shape[0], 0:image.shape[1]] = image

# Plotting
fig, ax = plt.subplots(figsize=(4.1, 4.1))
ax.imshow(combined_image, cmap='gray', extent=[0, main_img_w, main_img_h, 0])
plt.tick_params(top='off', bottom='off', left='off', right='off', direction="in", color='gray')
plt.xlim(0, main_img_w)
plt.ylim(main_img_h, 0)

# Set y-ticks to start from -50 to 400
new_yticks = np.arange(-50, 406 + 1, 50)
plt.yticks(new_yticks + 50, new_yticks[::-1])

# Draw edges and shortest paths
draw_edge_lines(ax, top_edge_points, bottom_edge_points)
shortest_paths = draw_shortest_distance_lines(ax, top_edge_points, bottom_edge_points)
center_points = draw_center_points(ax, shortest_paths)

# Draw tangents on the new center points
draw_tangents_only(ax, center_points)

# Draw the centerline curve
# Draw the centerline curve
draw_centerline_curve(ax, center_points)
draw_edge_lines(ax, top_edge_points, bottom_edge_points)
draw_tangents_only(ax, center_points)


# Define the vertical line position
vertical_line_position = 372  # Adjust this value as needed

# Define the horizontal line position
horizontal_line_position = 400  # Adjust this value as needed

# Define the x_position for draw_vertical_line1 function
x_position = 30  # Adjust this value as needed

# Draw the vertical line at a specified position
draw_vertical_line(ax, vertical_line_position, color='gray', linestyle='-', linewidth=1)

# Draw the horizontal line at a specified position
draw_horizontal_line(ax, horizontal_line_position, color='gray', linestyle='-', linewidth=1)

# Manually specify the arc center and theta position
arc_center = (359.5, 262)  # Adjust these coordinates as needed
theta_position = 90      # Adjust this value as needed
draw_arc_and_theta(ax, arc_center, theta_position, radius=25, theta_offset=10, arc_color='gray', theta_color='gray', arc_rotation=180)


# Save the plot as a PDF
plt.savefig('Final_Accurate_Centerline_and_Tangent_arc.pdf', format='pdf', dpi=1000)

# Show the plot
plt.show()
