## Supplementary material for "Humidity-driven shape morphing enhances fog harvesting in porous cactus spines": Software code S3

# Spine 2 with color code 

import cv2
import numpy as np
import matplotlib.pyplot as plt
import os
from scipy.optimize import curve_fit
from scipy.interpolate import UnivariateSpline
from matplotlib import colors

# Edge detection function (remains unchanged)
def detect_edges(image_path):
    image = cv2.imread(image_path, cv2.IMREAD_GRAYSCALE)
    blurred_image = cv2.GaussianBlur(image, (5, 5), 0)
    sobelx = cv2.Sobel(blurred_image, cv2.CV_64F, 1, 0, ksize=5)
    sobely = cv2.Sobel(blurred_image, cv2.CV_64F, 0, 1, ksize=5)
    gradient_magnitude = np.sqrt(sobelx**2 + sobely**2)
    gradient_magnitude = np.uint8(255 * gradient_magnitude / np.max(gradient_magnitude))
    _, binary_edges = cv2.threshold(gradient_magnitude, 100, 255, cv2.THRESH_BINARY)
    
    top_edge_points = []

    for x in range(binary_edges.shape[1]):
        for y in range(binary_edges.shape[0]):
            if binary_edges[y, x] > 0:
                top_edge_points.append([x, y])
                break
    
    return image, np.array(top_edge_points)

# New fitting function (simplified model)
def fit_function(x, A1, A2, A3, A4, A5):
    return A1 - A2 * np.exp(-A3 * x) + A4 * x**4 + A5 * x

# Updated fitting function to use curve_fit
def fit_weighted_function(top_edge_points):
    x = top_edge_points[:, 0]
    y = top_edge_points[:, 1]
    
    # Use curve_fit to find the best fit parameters for the new model
    initial_params = [10**-6, 10**-6, 10**-7, 10**-5, 10**-8]
    param_bounds = ([0, 0, 0, 0, 0], [np.inf, np.inf, np.inf, np.inf, np.inf])
    
    # Increase maxfev to allow more iterations
    fitted_params, _ = curve_fit(fit_function, x, y, p0=initial_params, bounds=param_bounds, maxfev=5000)
    
    return fitted_params

# Calculate the y-values using the new fitted function
def calculate_y_vals(fitted_params, x_vals):
    return fit_function(x_vals, *fitted_params)

# Calculate the tangent angles with Y-axis from the fit equation
def calculate_tangent_angles(fitted_params, x_vals):
    A1, A2, A3, A4, A5 = fitted_params
    dydx = A2 * A3 * np.exp(-A3 * x_vals) + 4 * A4 * x_vals**3 + A5
    angles = np.degrees(np.arctan(1 / dydx))
    return angles

# Process images, fit the new function, and plot the results
def plot_angles_with_radius(folder_path):
    color_map = plt.cm.viridis  # Use viridis color map
    tangent_angles_data = {}  # Dictionary to store tangent angles for each image
    
    # Get list of files and sort them numerically
    files = [f for f in os.listdir(folder_path) if f.endswith('.jpg') or f.endswith('.png')]
    files.sort(key=lambda f: int(os.path.splitext(f)[0]))
    
    # Create a single plot for all centerlines
    plt.figure(figsize=(4.1, 4.1))
    
    for idx, filename in enumerate(files):
        image_path = os.path.join(folder_path, filename)
        image, top_edge_points = detect_edges(image_path)
        
        # Invert the y-coordinates of the edge points for correct plotting
        top_edge_points[:, 1] = -top_edge_points[:, 1] + np.max(top_edge_points[:, 1])

        # Shift all y-values to make the first point's y-coordinate zero
        top_edge_points[:, 1] -= top_edge_points[0, 1]

        # Normalize x-values of edge points for plotting
        edge_x_vals_normalized = (top_edge_points[:, 0] - np.min(top_edge_points[:, 0])) / (np.max(top_edge_points[:, 0]) - np.min(top_edge_points[:, 0]))
        
        # Fit the new model to the remaining points
        fitted_params = fit_weighted_function(top_edge_points)
        
        # Generate 500 equally spaced points along the x-axis of the fitted function
        x_min, x_max = np.min(top_edge_points[:, 0]), np.max(top_edge_points[:, 0])
        x_vals = np.linspace(x_min, x_max - 1, 500)
        
        # Calculate the y-values using the fitted function
        y_vals = calculate_y_vals(fitted_params, x_vals)
        
        # Normalize x-values to the range [0, 1] for fitted line
        x_vals_normalized = (x_vals - np.min(x_vals)) / (np.max(x_vals) - np.min(x_vals))
        
        # Calculate the tangent angles using the fit equation
        tangent_angles = calculate_tangent_angles(fitted_params, x_vals)
        
        # Store the tangent angles data
        tangent_angles_data[filename] = tangent_angles
        
        # Plot the original edge points with normalized x-values
        plt.scatter(edge_x_vals_normalized, top_edge_points[:, 1], label='Edge Points', color='gray', s=10)
        # Plot the centerline with normalized x-axis
        plt.plot(x_vals_normalized, y_vals, label=f'Fit - {filename}', color=color_map(0.2 + 0.6 * (idx / len(files))))
    
    plt.xlabel('Normalized X')
    plt.ylabel('Y')
    plt.xlim(0, 1)
    plt.ylim(0, 400)
#     plt.savefig('fit_normalizedSpine2.pdf', bbox_inches="tight", transparent=True)
    plt.show()

    # Plot all tangent angles with the Y-axis in a single plot
    plt.figure(figsize=(5.1, 4.1))
    plt.tick_params(top='off', bottom='off', left='off', right='off', direction="in")
    for idx, (filename, angles) in enumerate(tangent_angles_data.items()):
        x = np.linspace(0, 1, len(angles))  # Normalize the x-values
        plt.plot(x, angles, label=f'{filename}', linestyle='-', color=color_map(0.2 + 0.6 * (idx / len(files))))
    
    plt.xlabel('Normalized Length, $s/L$')
    plt.ylabel('$ \\Theta (\\degree)$')
    plt.xlim(0, 1)
    plt.ylim(20, 100)

    # Adding the color bar with the truncated colormap
    new_cmap = colors.LinearSegmentedColormap.from_list('truncated_viridis', color_map(np.linspace(0.2, 0.8, 256)))
    sm = plt.cm.ScalarMappable(cmap=new_cmap, norm=plt.Normalize(vmin=1, vmax=300))
    cbar = plt.colorbar(sm, ticks=[1, 150, 300])
    cbar.set_label('Time (s)', rotation=-90, labelpad=20)

    plt.savefig('Spine2 Theta_vs_Normalized.pdf', bbox_inches="tight", transparent=True)
    plt.show()

# Define the input folder path
input_folder = '/Users/ovee/Desktop/Image/Jessica Data/Video_Spine2/Cropped/Test/mirror'
# input_folder = '/Users/ovee/Desktop/Image/Jessica Data/Videos_Spine1/S1_A1_R6/Cropped/1/mirror'

# Process images and draw the radii, tangents, and edge points with the new fitting function
plot_angles_with_radius(input_folder)
